# A generative learning model for saccade adaptation

**DOI:** 10.1101/492512

**Authors:** Carlos R. Cassanello, Florian Ostendorf, Martin Rolfs

**Affiliations:** Department of Psychology, Humboldt Universität zu Berlin, Philippstr. 13 Haus 6, 10115 Berlin, Germany; Department of Bernstein Center for Computational Neuroscience Humboldt Universität zu Berlin, Philippstr. 13 Haus 6, 10115 Berlin, Germany; Department of Neurology Charité – University Medicine Berlin, Augustenburger Platz 1, 13355 Berlin, Germany

**Keywords:** visually-guided saccades, sensorimotor learning, oculomotor plasticity, parameter estimation

## Abstract

Plasticity in the oculomotor system ensures that saccadic eye movements reliably meet their visual goals—to bring regions of interest into foveal, high-acuity vision. Here, we present a comprehensive description of sensorimotor learning in saccades. We induced continuous adaptation of saccade amplitudes using a double-step paradigm, in which participants saccade to a peripheral target stimulus, which then undergoes a surreptitious, intra-saccadic shift (ISS) as the eyes are in flight. In our experiments, the ISS followed a systematic variation, increasing or decreasing from one saccade to the next as a sinusoidal function of the trial number. Over a large range of frequencies, we confirm that adaptation gain shows (1) a periodic response, reflecting the frequency of the ISS with a delay of a number of trials, and (2) a simultaneous drift towards lower saccade gains. We then show that state-space-based linear time-invariant systems (LTIS) represent suitable generative models for this evolution of saccade gain over time. This *state-equation* algorithm computes the prediction of an internal (or hidden state-) variable by learning from recent feedback errors, and it can be compared to experimentally observed adaptation gain. The algorithm also includes a forgetting rate that quantifies per-trial leaks in the adaptation gain, as well as a systematic, non-error-based bias. Finally, we study how the parameters of the generative models depend on features of the ISS. Driven by a sinusoidal disturbance, the state-equation admits an exact analytical solution that expresses the parameters of the phenomenological description as functions of those of the generative model. Together with statistical model selection criteria, we use these correspondences to characterize and refine the structure of compatible state-equation models. We discuss the relation of these findings to established results and suggest that they may guide further design of experimental research across domains of sensorimotor adaptation.

**Author Summary:** Constant adjustments of saccade metrics maintain oculomotor accuracy under changing environments. This error-driven learning can be induced experimentally by manipulating the targeting error of eye movements. Here, we investigate oculomotor learning in healthy participants in response to a sinusoidally evolving error. We then fit a class of generative models to the observed dynamics of oculomotor adaptation under this new learning regime. Formal model comparison suggests a richer model parameterization for such a sinusoidal error variation than proposed so far in the context of classical, step-like disturbances. We identify and fit the parameters of a generative model as underlying those of a phenomenological description of adaptation dynamics and provide an explicit link of this generative model to more established state equations for motor learning. The joint use of the sinusoidal adaption regime and consecutive model fit may provide a powerful approach to assess interindividual differences in adaptation across healthy individuals and to evaluate changes in learning dynamics in altered brain states, such as sustained by injuries, diseases, or aging.

## Introduction

The accuracy of saccadic eye movement is maintained through mechanisms of saccade adaptation, which adjust the amplitude [1–3] or direction [4–6] of subsequent movements in response to targeting errors. As online visual feedback cannot be used to correct the ongoing movement, saccadic eye movements need to be preprogrammed and adaptation must largely rely on past experience and active predictions [7,8] rather than closed-loop sensory information.

To induce saccade adaptation in the laboratory [1], participants are instructed to follow a step of a target stimulus with their eyes and this visual cue is then displaced further during the saccade eye movement. Typically, this second, intra-saccadic step (ISS) is constant across trials and directed along the initial target vector towards smaller or larger saccade amplitudes. Although the ISS is visually imperceptible [9], saccades adjust their amplitude to compensate for the induced error. In phenomenological analyses of such saccade adaptation data, the amount of adaptation is usually quantified by comparing saccade gain values before and after the adapting block and interpolating an exponential fit in between [1–3,10].

We recently presented a version of this paradigm in which the ISS (the disturbance responsible for inducing adaptation) follows a sinusoidal variation as a function of trial number ([11,12]; see also [4,13,14]). We reported that gain changes were well described by a parametric functional form consisting of two additive components. One component was a *periodic response* reflecting the frequency of the ISS that was adequately fitted with a lagged but otherwise undistorted sinusoid. The second component constituted a *drift of the baseline* toward lower saccade gain (larger hypometria) that was appropriately accounted for using an exponential dependence.

Here, we investigate whether a generative algorithm that models saccade gain modifications on a trial-by-trial basis by learning from errors made on previous trials can account for this response. To this end, we implemented and fit a series of state-space models in which a modified delta-rule algorithm updates a hidden or latent variable (for which the experimentally observed adaptation gain is a proxy) by weighting the last experienced visual error, in addition to other error-based and non-error based learning components [8,15–23].

We adopt the approach that these algorithms are linear time-invariant systems (LTIS), in that their coefficients are time and trial-independent. LTIS models, also known as linear dynamical systems (LDS) have been successfully used in a number of motor adaptation studies [8,19–22,24–27]. Applied to saccade adaptation, they may predict the dynamics of the saccade amplitude itself as well as various forms of movement gain typically used in describing adaptation [2,3,10,11]. Our first goal was to establish empirically whether LTIS models could fit the data recorded with a sinusoidal adaptation paradigm, as efficiently as when using a constant (fixed) ISS. Once we have established this point, we will explore the relation between the predicted phenomenological parameters [11,12] and the learning parameters of the underlying generative model, as well as their potential dependence on the perturbation dynamics.

We first analyze the ability of a family of generative models to describe experimental recordings of saccade adaptation by fitting the relevant learning parameters. We then perform statistical model-selection analysis to determine those that best fitted the same data in the various experimental conditions. We fitted models to two data sets, a previously published one [11] and a variation of that paradigm that extended the range of frequencies of the sinusoidal variation of the ISS. Both data sets contrasted two established saccadic adaptation protocols [11,12]: Two-way adaptation (i.e., bidirectional adaptation along the saccade vector of saccades executed along the horizontal meridian) and Global adaptation (i.e., adaptation along the saccade vector of saccades executed in random directions). We then explore consequences for current models of motor learning and suggest possible modifications that may be required to generate a suitable description of sensorimotor learning during sinusoidal saccadic adaptation. In conducting this selection, we confirm that a single learning parameter model (a state-equation with just an error-based learning term; cf. [19]) does not suffice to fit the data. We then demonstrate that including an extra term that weights the next-to-last trial s error provides a better fit for the Two-way type of adaptation.

This learning rate has the intriguing feature that it has negative values for all frequencies, suggesting an active unlearning of the next-to-last trial s feedback error, close, but not equal in magnitude to the learning rate of the last trial’s error. We discuss possible functional roles of these processes for oculomotor adaptation in natural situations, where saccadic accuracy is expected to exhibit slow dynamic changes across time.

## Methods

### Procedure

We re-analyzed the data we recently collected using a fast-paced saccade adaptation paradigm with a sinusoidal disturbance. We had previously described these data by fitting a phenomenological model that we identified using statistical model selection. For details on the experimental procedures pertaining to this original data set (henceforth, ORIG) and to the selection of the functional form of this phenomenological model, please refer to our former communication [11].

We applied the same experimental procedure in collecting further data with an enhanced range of frequencies. In this case, thirteen participants ran two sessions with similar Two-way and Global adaptation protocols as used in previous reports [11,12]. In short, Two-way adaptation refers to bidirectional adaptation along the saccade vector of saccades executed along the horizontal meridian. In turn, Global adaptation refers to adaptation along the saccade vector of saccades executed in random directions.

In collecting this dataset (henceforth, FREQ), each session had 2370 trials divided in 11 blocks. Odd numbered blocks had 75 no-adaptation trials (zero ISS). The five even-numbered blocks consisted of 384 trials each with a sinusoidal disturbance similar to that used before but with frequencies of 1, 3, 6, 12 and 24 cycles per block (i.e., 384, 128, 64, 32, and 16 saccades per cycle, respectively). The order of adaptation blocks was randomly interleaved for each observer and type of adaptation. The program was paused after each adaptation block, giving participants some resting time, and we calibrated eye position routinely at the beginning of each non-adapting (odd-numbered) block. In each trial, the pre-saccadic target step was fixed at 8 degrees of visual angle (dva). The subsequent second step (ISS) then ranged between −25% and +25% of the first step, changing size according to a sine function of trial number.

The Ethics Committee of the German Society for Psychology (DGPs) approved our protocols. We obtained written informed consent from all participants prior to their inclusion in the study. The present study conformed to the Declaration of Helsinki (2008).

### Data analysis and phenomenological model

#### Modeling of the saccadic response

In a double-step adaptation paradigm [1], after a fixation interval the fixation target *FP*(*n*) undergoes a first step to become the target of a saccade, displayed at the pre-saccadic location *TP*1(*n*). Because the eyes might have been stationed at a location *EP*1(*n*) close to but different than *FP*(*n*), we define the pre-saccadic target amplitude *preTP*(*n*) = *TP*1(*n*) – *EP*1(*n*), with origin at *EP*1(*n*) rather than *FP*(*n*) and keep this convention throughout the study.

The second step of the McLaughlin paradigm (i.e., the target displacement inducing a feedback error) then shifts the target during the saccade to a position *TP*2(*n*)(so that *ISS*(*n*) = *TP*2(*n*) – *TP*1(*n*). Therefore, the post-saccadic target amplitude (at or immediately after saccade landing) is given by the identity: *postTP*(*n*) = *preTP*(*n*) + *ISS*(*n*). For convenience, we will define a *target gain, t*(*n*), as the ratio of the post-saccadic target amplitude to the pre-saccadic one, as well as a *disturbance gain, d*(*n*), as the ratio of the second to the first target steps, i.e., the ratio of the *ISS* to the saccade proxy:

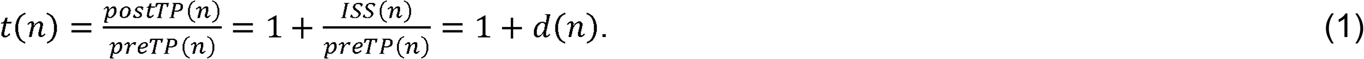

In the general case, there would be a constant and a variable component in the second target step, *ISS*(*n*) = *C* + *V*(*n*). In our sinusoidal adaptation paradigms, *C* = 0 and 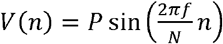 is a sine function of the trial number so that:

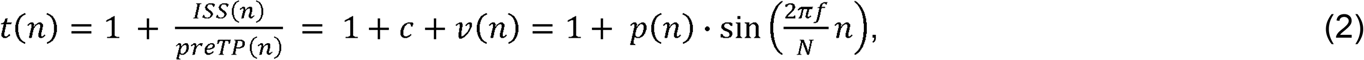

where *c* and *v*(*n*) are the ratios of the constant and variable part of the *ISS* to the pre-saccadic target amplitude. In the sinusoidal paradigms, *f* is the frequency of the sinusoid in cycles per block, *N* is the number of trials in an adaptation block, and *n* is the index of the current trial. At fixed amplitude, the dynamics of the disturbance is fully determined by its angular frequency 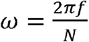 that characterizes the rate of change of the sinusoid in each trial. *P* is the maximum absolute magnitude of the variable part *V*(*n*), i.e., the ‘amplitude’ of the sinusoid that defines the *ISS*. It was fixed at 2 dva throughout all sinusoidal adaptation datasets. Therefore, *ISS*(*n*) changed in magnitude periodically and in a sinusoidal fashion between approximately −25% and +25% of the magnitude of the pre-saccadic target eccentricity (*preTP*(*n*)), which was held approximately fixed at 8 dva in all datasets. Finally, *p*(*n*) is the ratio of *P* and *preTP*(*n*), and had an approximately constant value of 0.25 across the sinusoidal datasets (the slight dependence on the trial number was a consequence of the slight dependence of the normalizing factor *preTP*(*n*) on the trial number; in actuality, the magnitude held constant at 8 dva across the experiment was *TP*1(*n*) – *FP*(*n*), which differed slightly but not systematically from *TP*1(*n*) – *EP*1(*n*)). Given that we used integer number of cycles across all sinusoidal adaptation experiments, we expressed the frequency in cycles per block (cpb). We set the initial phase to zero, which means that the magnitude of the ISS starts at zero in the direction of positive *ISS* (outward second-steps of the saccade target) first. **Equation 2** provides a complete description of the stimulus that we used. Yet, for the analyses pursued here and to make closer contact with our phenomenological characterization of oculomotor responses in sinusoidal adaptation [11], we will further define a *stimulus gain, s*(*n*), to be the disturbance gain normalized to (i.e., divided by) its maximum absolute value. Therefore, *s*(*n*) would range within ±1 in units of its maximum amplitude following a sinusoidal variation with trial number:

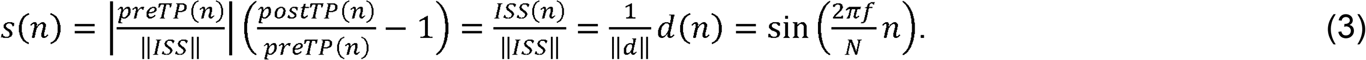

Saccade amplitude adaptation is usually described in terms of the changes in *saccade gain* (*SG*(*n*)), defined as the ratio of the saccade amplitude (*SA*(*n*)) to the pre-saccadic position error (*preTP*(*n*)). During non-adapting trials and at the beginning of the adaptation blocks, *SG*(*n*) is typically slightly smaller than 1, which means that the saccade undershoots the target. Since we are interested in keeping track of the excursions of the saccade gain with respect to a perfect completion of the saccade that matches *preTP*(*n*) exactly, we shall define an *adaptation gain* subtracting one from the usual saccade gain and normalized to the maximum absolute value of the *ISS*,

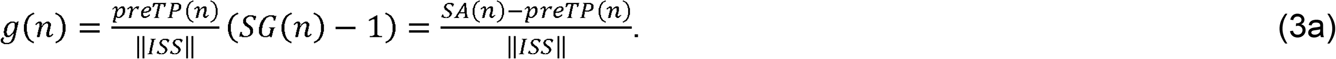

The adaptation gain represents the residual of the saccade gain with respect to perfect landing. When a saccade lands exactly on the first target step (a perfectly accurate saccade), the saccade gain will be one while the adaptation gain will be zero. Therefore, the adaptation gain uses perfect landing as the origin of coordinates and quantifies departures from this ideal goal state. Clearly, in both descriptions the reference represents a state of no adaptation. The adaptation gain description may be viewed as following the evolution of the error rather than that of the full eye movement. As long as the true underlying learning model is strictly linear, both descriptions must be equivalent since they relate to each other by a shift. We used the adaptation gain, *g*(*n*), in our previous reports [11,12] to provide phenomenological parametric description of sinusoidal adaptation data and it is also commonly used within motor control research. Throughout the manuscript we shall use *SG*(*n*) or *g*(*n*) as the relevant behavioral variables describing the data, which are computed directly from the experimental measurements of the eye and target positions in each trial.

#### Assessment of the evidence in favor of a model

In implementing the phenomenological parameter estimation, we adopted a Gaussian likelihood for the data given the model. This likelihood can be maximized with respect to the parameters at a fixed but unknown width. Instead we adopted the following procedure [11,13]. Using Bayes theorem, priors for the parameters to be estimated, and assuming a constant prior probability for the data, we can obtain a joint probability amplitude for all parameters that can be marginalized to extract individual probability amplitudes for each parameter. In this process, the width of the Gaussian likelihood is a nuisance parameter that we integrate out using a non-informative prior [13,28,29]. Once such integration is conducted, the volume of the resulting probability density (given the data) provides an estimate of the odds that the model would provide a reasonable description of the data. Here we provide a full model consisting of six parameters (sinusoidal entraining of the oculomotor response riding over a baseline drift) that we want to compare to a partial model (the drift of the baseline alone) and to a minimal model consisting of the mean of the adapting block with variance equal to the variance of the recorded data over that block. To establish which situation is more likely across different number of parameters, we take the log of the ratio of the odds across the models. The resulting magnitude is the *evidence* that the data are in favor of a particular model and is measured in *decibels* (db). When this magnitude is positive, the odds favor the model in the numerator, with evidence higher than 3 db indicating that this model is significantly favored to the one in the denominator. We use this metric to assess the quality of our parameter estimation.

#### Statistics

Throughout the manuscript we report results as mean ± SD for individual data and mean ± SEM when we discuss group data. In the phenomenological fittings, to determine average parameters from the parameter estimation other than the frequency, we computed the mean and variance for each parameter and participant as the first two moments of the corresponding posterior probability distribution and took the average of the means weighted by their standard deviations (square root of the estimated variance) to generate each point on the population plot. Alternative estimators (e.g., the modes of the posterior distributions, with and without weighting) gave qualitatively similar results.

### Modeling of the sensorimotor learning process: the modified delta-rule state equation

To investigate generative models, we adopt the following rationale. In each trial, the oculomotor system must generate a motor command to produce the impending saccade. This needs to be calibrated against the actual physical size of the required movement [15,20,22,24,30,31].

If the saccade fails to land on target, the motor command needs to be recalibrated based on preexisting calibrations, and we will hypothesize that those changes take place in an obligatory manner (cf. [19]) through additive, error-based modifications attempting to ameliorate post-saccadic mismatches between the eyes landing position and target location.

We model the underlying sensorimotor learning using linear time-invariant systems (LTIS) because the model parameters (or the learning coefficients) are time independent in each experimental block, although they can vary across experimental conditions or phases [32]. These models are closely related to linear dynamical systems (LDS; cf. [20–22]), except that here we only address noise-free models.

Because saccades are extremely rapid movements that do not admit reprogramming in mid-flight, it is assumed that all gain changes take place in between saccades. In our models, therefore, the error-based correction terms weight errors that were experienced in previous saccades. As a consequence, in the estimation of the forthcoming event, the post-saccadic stimulus gain is not compared against the adaptation gain measured for that trial but against the previous estimate of the gain. To justify these assumptions, it is usually assumed that the motor system sends an *efference copy* of the motor command to the sensory areas, which enables prediction of the sensory consequences of the movement and therefore avails comparison to experienced post-saccadic feedback [7,19,20,31,33–35].

We will assume that the values of saccade and adaptation gains observed and extracted from the recorded data (i.e., 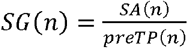 and 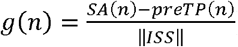) are adequate proxies of that motor calibration process. Yet the calibration itself is an internal feature of the brain and therefore the *adaptation gain* that enters the generative algorithm (the state-equation) that we intend to study is a *hidden variable* representing the *internal state* of the system. A model providing its temporal evolution can then be fitted to the data; yet the variable itself is not experimentally accessible. We denote the internal variable associated to the saccade gain by *z*(*n*). To describe the evolution of this *state* variable we introduce the state-equation:

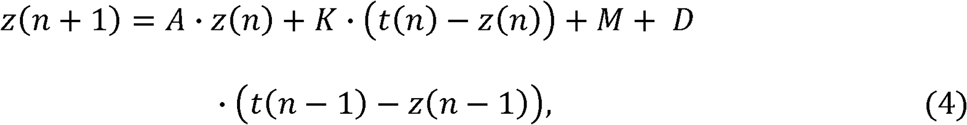

supplemented with an initial condition that sets the initial value *z*(1) = *G · t*(1). Here, the target gain *t*(*n*) is available from recordings in each trial and we shall assess how well the *prediction* of the saccade gain (*z*(*n* + 1), provided by **Equation 4**) fits the recorded data *SG*(*n*). The first term on the RHS of the equation is a persistence term. The persistence rate *A* determines how much of the estimate of the state variable at trial *n* is transferred to the estimate at the next trial [8,25,36]. Therefore, its magnitude is expected to be typically slightly smaller than 1 and it is set to be 1 in the models that do not include its effect. The second term weights the discrepancy between the gain of the target at trial *n* and the predicted gain of the movement under the underlying assumption that the size of the state variable is an adequate proxy for the (sensory) consequences of the movement. The weighting coefficient *K* is called *learning rate. M* embodies any systematic effect (drift or bias) that takes place in each trial but is not directly determined by the sensory feedback 2[37]; we shall call it a drift parameter. The last term is a second error-based correction term that weights the discrepancy between the gain of the target and the estimate of the movement at a trial other than the last error with and additional (distal) learning rate *D*. For concreteness we shall assume that this correction is based on the sensory feedback arising from the next-to-last trial. However, we shall return to this specific assumption further in the **Discussion**. Note that with the inclusion of this hypothetical double error sampling the full model of **Equation 4** (and **Equation 5** below) becomes an algorithm that coherently uses two delayed feedbacks to estimate the state of a single internal variable that models the sensory consequences of the intended motion.

#### Formatting of the data for fittings of the learning model

To be able to consistently compare results from this manuscript with the phenomenological analyses of the data presented in our earlier report, we will write the generative model in terms of a state variable associated to the *adaptation gain* of **Equation 3a** (cf. [11], and therefore naturally defined as 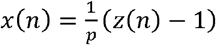. Applying these changes, we obtain:

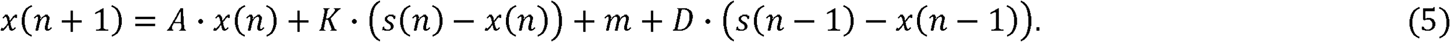

As suggested by **Equations 4** and **5**, a sensorimotor learning model can be written in terms of hidden variables that would be naturally associated with the saccade gain or the adaptation gain defined in **Equations 3** and **3a** respectively. When transitioning from the saccade gain to the adaptation gain description in this linear model, the only parameter of **Equation 4** susceptible to changes is *M*, which we indicated in **Equation 5** using the lower-case *m* instead. Throughout the manuscript we adopt the adaptation gain (defined above), as the state variable to characterize the internal model and **Equation 5** as its relevant state-equation. In this description, the stimulus gain reduces to a pure sinusoidal disturbance with zero mean (i.e., with no static component), which minimizes confounds between the effects of the retention rate *A* and the drift parameter *m*.

Because movement gains are computed from experimental observations, models of motor control often include a second equation that maps the estimates of the hypothesized internal variable to real-world observations (see, e.g., [20,21]). In our simplified analyses and again invoking the pre-programmed nature and accuracy of saccades, we set this second (observation) stage to be an identity.

### Estimation of the learning parameters, model classification and model selection

We conducted our analyses using the full form of **Equation 5**. We were interested in determining which model suffices to account for the data with the least number of parameters. The magnitude being learned is *x*, the internal representation of the adaptation gain of the imminent saccade. This gain has value zero upon the ideal outcome of perfect movement accuracy and in that respect, it can be interpreted as the gain of an internal prediction error. Using **Equation 5**, we generated the predicted values of *x*(*n*) in each condition and for each participant, and then fitted a number of models that differed from each other in which parameters were estimated. When a parameter among *K, m*, or *D* was not present, the corresponding term was removed from **Equation 5**. Note however, that when the parameter was not included as a fitting parameter, its value was set to unity (i.e., *A* = 1). In the case of the initial value *G*, we obtained an estimate by taking the average of the first five valued of the gain. We proceeded in this way because the initial value of the state of the system is unknown and, while the first recorded value of the gain could be considered a proxy for such initial state, execution and motor errors could yield a value of the gain significantly different than the actual initial state of the system; we averaged over 5 trials to alleviate this problem. In models where the initial value of the gain was left free to become a fitting parameter, this average over the first five saccades was used as an initial value for the fitting routine for that particular parameter. Improvements can be achieved by letting the initial condition become an extra parameter. We discuss below the interpretation of using the initial condition as a fitting parameter of the model.

In view of these features of the generative model, a natural classification of the models tested arises as follows: given the parameters *K,A,m, D*_1_,⋯,*D_w_,G*, we will 1) include *K* in every model because we are modeling intrinsic learning where we assume that learning from the last experienced feedback is always present as well as obligatory [19,20,22,31]; 2) models will be generated by adding successively the parameters *A, m*, and *D*, of which one or more could be present but in this study we restrict ourselves to learning possibly from only one extra feedback in the past; 3) *G* is an optional parameter that is included in an attempt to alleviate extreme effects of the initial condition(s) as explained above. By applying points 1) through 3), sixteen different models can be generated. For reasons to become clear below we would group them in four families according to whether or not they contain the bias term (*m*) and the additional error term (with learning rate *D*): only (although with *A* = 1 when omitted) *KA, KG, KAG* feature zero bias and a single error term; *Km, KAm, KmG, KAmG* are models with a single error term that allow bias; *KD, KAD, KDG, KADG* have no bias term but sample two errors, and *KmD, KAmD, KmD* G, *KAmDG* feature both a bias term and learn based on double error sampling. Therefore, the simplest model had a single fitting parameter (the learning rate *K*, cf. [19]) and was obtained by setting *A* = 1, removing the terms that involved *m* and *D*, and setting the initial value *G* to be the mean of the first five values of the gain in the block. The full model had all five as fitting parameters.

All parameters of the generative models were estimated by fitting the model to the experimental data using MATLAB function nlinfit; 95% parameters confidence intervals were computed using MATLAB function nlparci and predicted response for the hidden variable *x* with its corresponding 95% confidence intervals were obtained from MATLAB function nlpredci.

All 16 models were fitted to each individual participants’ data, parameters were extracted for each model, and models were compared using the Akaike information criterion (AIC; [38–41]) by computing Akaike weights across models for each participant. Finally, these weights were averaged across participants for each model in each condition.

### Using the generative model to predict the parameters of the phenomenological description of the adaptation gain

The adaptation gain of the oculomotor response to a sinusoidal disturbance is best described by a phenomenological function consisting of a decaying exponential added to a lagged but otherwise undistorted sinusoid [11]. The sinusoidal component of the response onsets at the beginning of the adaptation block but all fittings include the pre-adaptation block as well. The frequency of the stimulus disturbance is matched closely by the gain. To fully describe the response, five extra phenomenological parameters are required: amplitude (*a*) and lag (*ϕ*) of the periodic part of the error gain complete the description of the periodic part. The exponential decaying component that describes the baseline on which the periodic response rides requires other three: an asymptotic value (*B*_0_) where the baseline stabilizes at large trial number, a timescale (*λ*) in which the baseline reaches 1/*e* of the full decay, and the amplitude of the decay (*B*):

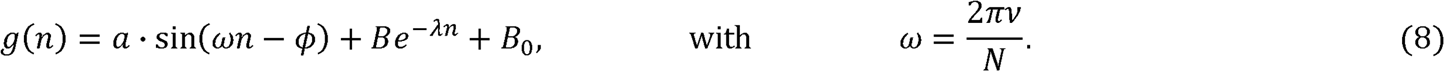

We use here the same denominations used in our previous report [11], except for changing the name of the timescale to *λ* to prevent confusion with the amplitude of the periodic component *a*. To estimate parameters of the phenomenological functional form that best fits the data we used the same general procedure and parameter estimation algorithm implemented in our earlier contributions [11,13]. Solving the state-equation via iteration in the simpler case where the system learns only from the last experienced feedback (cf. **S1 Appendix**), or borrowing techniques from the theory of LTIS reveals a correspondence between these phenomenological parameters and the coefficients of the generative model of **Equation 5**. (A complete derivation of the phenomenological parameters as functions of the generative ones is not presented here due to space limitations; details about the analytical procedures adopted can be found in [42]). Depending on the parameters that each generative model includes, the functional form and value of the phenomenological coefficients may change. Here we are interested in assessing which theoretical prediction of the relation among phenomenological and generative model parameters matches the data best as a way to validate the underlying sensorimotor learning algorithm.

### Lag and amplitude of the periodic response

The lag of the periodic response of the error gain derived from the (full version of the) generative model of **Equation 5** including the next-to-last feedback-error term is given by:

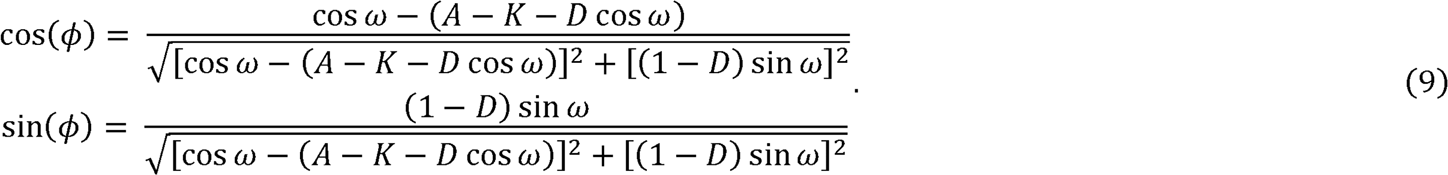

In models without next-to-last feedback term *D* should be set to zero; in models that do not have *A* as a fitting parameter, its value should be set to 1 in **Equation 9**.

The periodic component of the response to a sinusoidal disturbance in models where the next-to-last feedback is included can be written as:

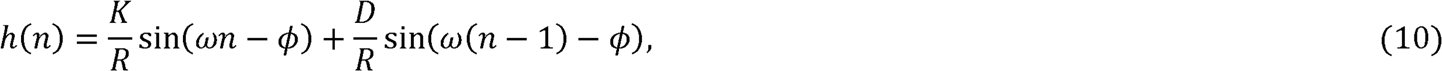

where 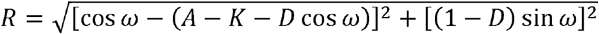.

**Equation 10** shows that if *D* = 0 we recover the solution expected by iteration when there is learning from the last error only. Then the amplitude of the periodic component (*a*) in **Equation 8** can be read out directly to be 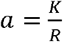. When *D* ≠ 0 we need to re-write **Equation 10** so that it matches the periodic part of **Equation 8**. After some algebra **Equation 10** can be recast as:

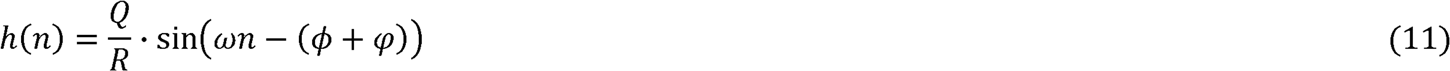

where

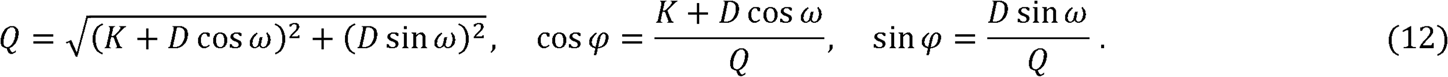

**Equations 9** to **12** clarify the effect of the presence of the next-to-last error learning rate *D*. **Equation 9** shows how the *bare* lag *ϕ* changes when *D* is present. Yet, it would be incorrect to compare the fitted values of the phenomenological lag to **Equation 9**. The reason is that the second contribution in **Equation 10** modifies not only the amplitude of the periodic component to the new value 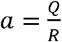, but it also adds the shift *φ* to the lag. Therefore, if there were also learning from the next-to-last error, the observed (behavioral) lag should be compared to *ϕ + φ*.

### Baseline drift parameters

Following a sinusoidal disturbance, the baseline of the error gain will approach an asymptote at large trial number that can be written as a function of parameters of **Equation 5** as (see also **S1 Appendix**):

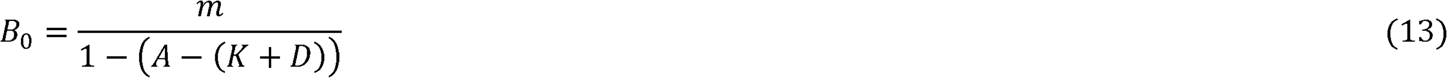

The timescale *λ* for the decay of the baseline, has units of 1/trials and it is defined by:

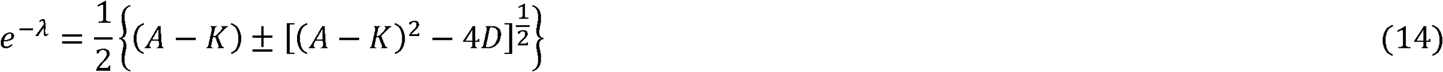

**Equation 14** provides the weights of the *impulse response* that generates the integral solution by convolving the stimulus (i.e., *s*(*n*); cf. **S1 Appendix**, [42]). The inverse of the timescale parameter *λ* gives the number of trials over which the stimulus is integrated. Beyond this *window of integration*, the weighting of the stimulus would have reduced enough to ignore further contributions. When *D* = 0, the integration weight becomes *e*^−*λ*^, which is positive and smaller than 1, provided that the learning rate *K* < 1 and *A*~1. When *D* ≠ 0, **Equation 14** provides timescales for two modes that compose the integral solution of the state-equation. These result from the addition or subtraction of the second term in braces. If the parameter *D* is negative, the second term inside the braces becomes slightly larger than the first. The timescale resulting from the addition is positive and can be expressed as a decaying exponential. The subtraction solution is negative and of small magnitude and, therefore, it will decay much faster when raised to the trial number. It introduces small additive fluctuations to the exponential decay of the addition solution without changing its overall behavior. Critically, diverse sizes of the learning parameters may result in smaller or larger timescales in models with *D* ≠ 0 compared to models where *D* = 0(cf. **Results** section and **S1 Appendix**).

To recap, **Equation 8** has four phenomenological parameters that we shall explore in further detail: *B*_0_, *λ*, *a*, and *ϕ*. The former two parameters are already familiar from phenomenological descriptions of data in paradigms using fixed-sized second-step for the target. The latter are new, arising in paradigms with sinusoidal disturbances.

The amplitude of the decay of the baseline also bears dependence on the learning rates as well as on the initial condition. Because of the strong influence of the initial condition on this parameter, we refrain from a comparison of the behavioral fittings to the predictions from the generative model for this case.

Part of the material discussed in this contribution have been presented in the form of posters or slide presentations [43,44].

## Results

### Analysis of the data at the phenomenological level

To obtain a general idea of patterns present in the data, we first collapsed the data for each stimulus frequency and adaptation type across participants (group data). We fit these data using a piecewise continuous function given by the addition of a monotonic (exponential) decay of the baseline -spanning both pre-adaptation and adapting trials- and a periodic entraining of the oculomotor response to the sinusoidal stimulus that begins at the onset of the adaptation block. This choice was supported by the fact that we had confirmed using statistical model selection criteria (i.e., AIC and BIC, [38–41,45]) that this functional dependence was the best descriptor of the oculomotor response among the set of models tested in Cassanello et al. [11]. For illustration purposes only, **Fig 1** shows the group data in each dataset, along with the fits resulting from the parameter estimation based on the phenomenological model of **Equation 8**. The same parameterization was used to fit each participant’s run. **Figs 2** and **3** summarize the estimation of the phenomenological parameters entering **Equation 8**. **Fig 2** shows the values of mean ± SEM of the parameters estimated from every individual dataset for each frequency and adaptation type.

**Fig 1.**
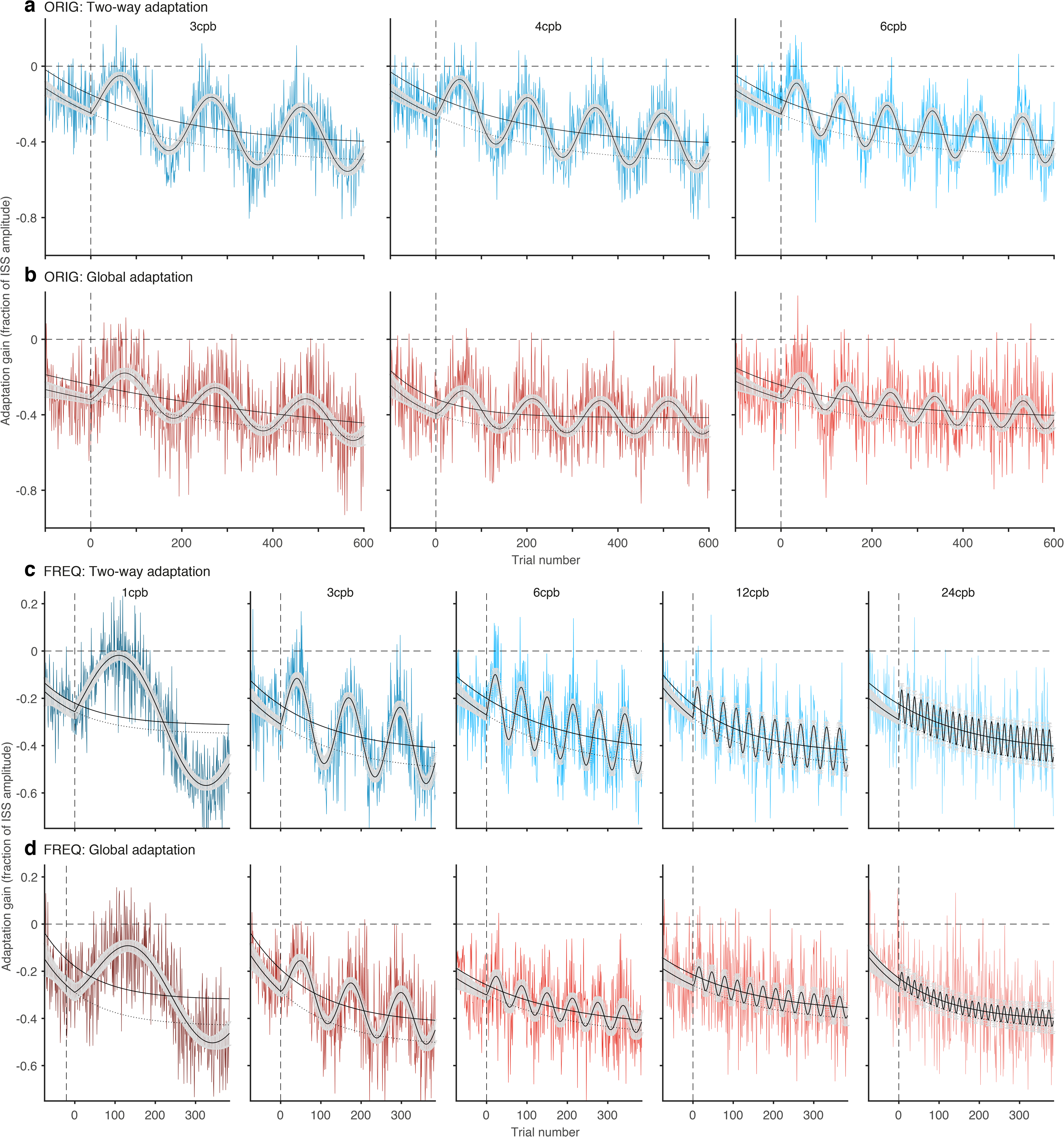
Fits of the phenomenological model to the experimental data. The plots show adaptation gain (colored lines) averaged over individuals in the (**a**) Two-way adaptation and (**b**) Global adaptation condition of the ORIG data set (reported in [11]), as well as the (**c**) Two-way adaptation and (**d**) Global adaptation condition of the FREQ data set, using the same paradigm over an extended range of frequencies. The fit (black line) is based on **Equation 8**. The same equation was fitted to data from each participant in each condition and experiment, to estimate phenomenological parameters on an individual bases. For illustration purposes only, the figure depicts fittings done over the averages along with 95% confidence intervals (gray shaded areas). The black dotted lines indicate the time evolution of the baseline if the amplitude of the periodic response were zero, corresponding to a drift only model. The solid black lines indicate the approximate middle-point locations of the periodic component.

**Fig 2.**
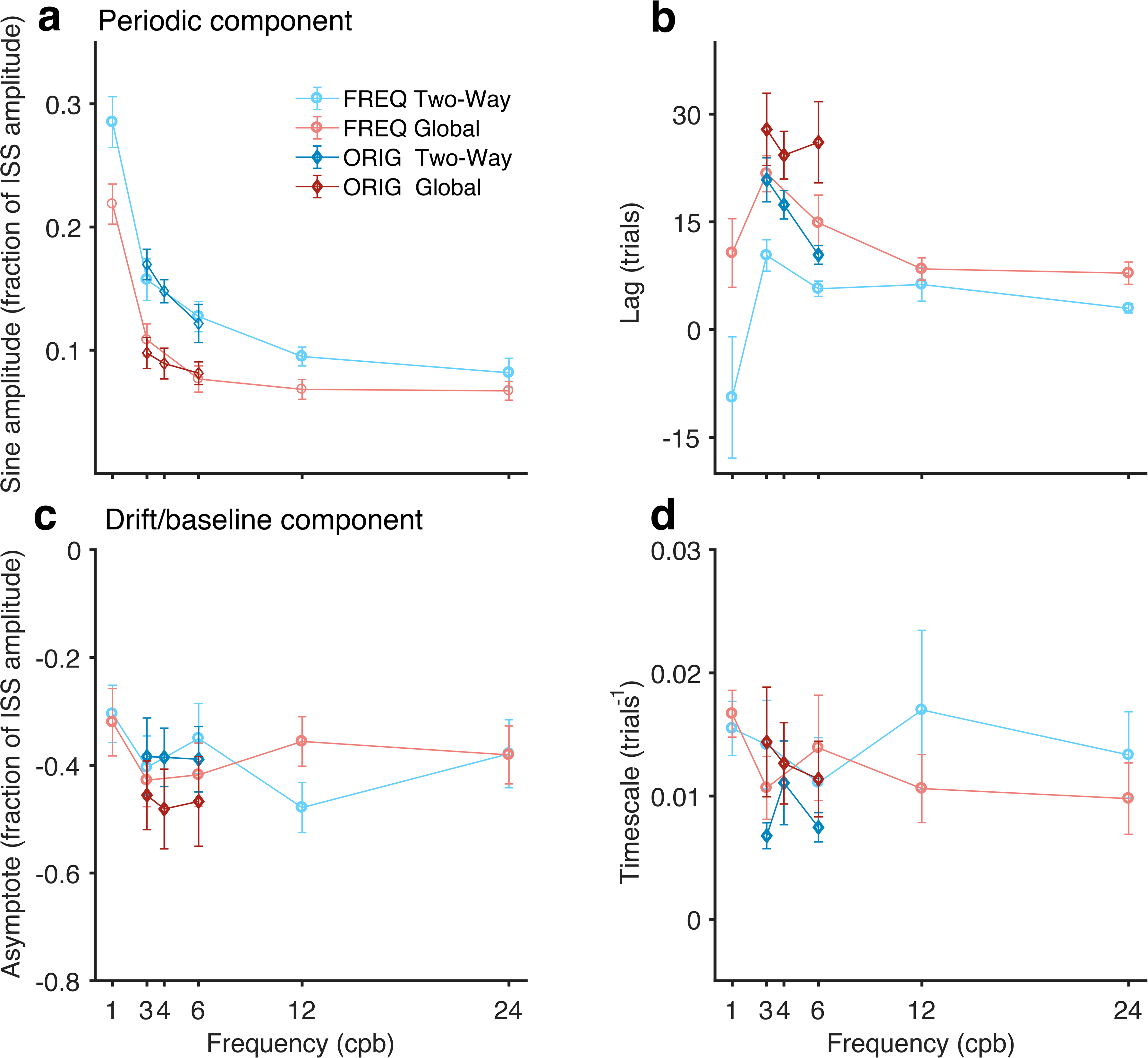
Phenomenological parameters as a function of ISS frequency, estimated from both datasets (ORIG, diamonds, and FREQ, circles). (**a,b**) Amplitude (**a**) and Lag (**b**) parameters of the periodic (sinusoidal) component of the response. (**c,d**) Asymptote (**c**) and timescale (**d**) parameters of the monotonic drift of the baseline toward greater hypometria. Each point is a condition defined by type of adaptation and ISS frequency. Blue and red colors correspond to horizontal Two-way and Global adaptation, respectively. Error bars are SEM across participants. These four parameters are further compared to the values predicted by the solution to the generative models tested.

Some features are readily apparent from these plots. First, the frequency of the ISS is reliably estimated (cf. **Fig 3b,d**). Second, the amplitude and the lag of the periodic components of the adaptation gain decay with increasing frequency of the stimulus (**Fig 2a,b**). The amplitudes of the periodic component are systematically larger in Two-way adaptation, while the lags observed in global adaptation are systematically larger than in the Two-way case. The systematic decay of the values of the lag with increasing frequency does not seem to extend to the smallest frequency (1 cpb in the new dataset). This may be related to the fact that at such low frequency the stimulus resembles more the behavior of a ramp that then turns rather than a truly periodic disturbance.

**Fig 3.**
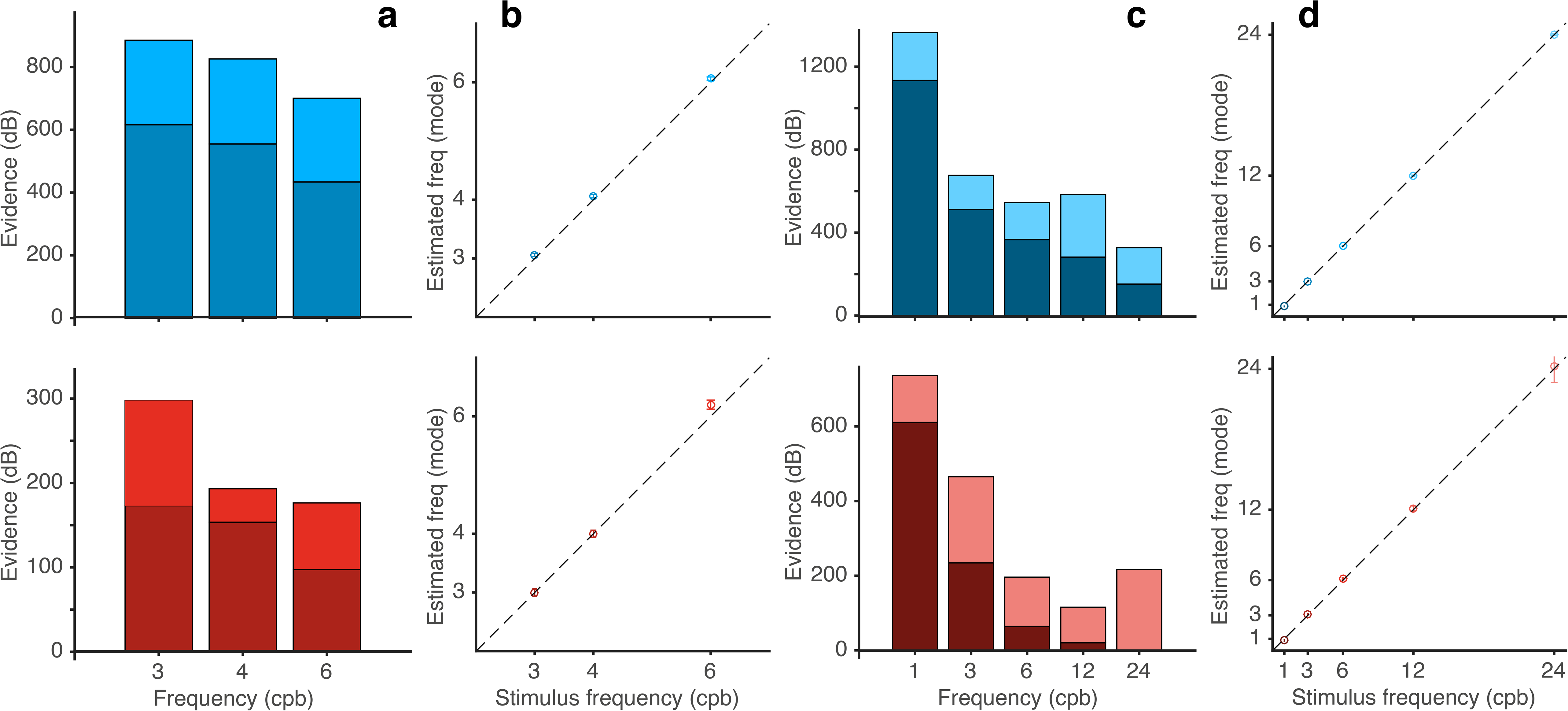
Assessment of the quality of the fits of the parametric phenomenological model to the group data. (**a,c**) Each bar is split into the log of the odds ratio of the full model to a drift only model that lacks the sinusoidal component (darker tone of the bars) added to the log of the odds ratio of the drift only model to the noise only model described above (lighter tone of the bars). For all but one condition (Global adaptation, 24 cpd), the full model provides the best account of the data. (**b,d**) Estimates of the frequency of the periodic component of the oculomotor response, for dataset ORIG (**b**) and dataset FREQ (**d**). Error bars are SEM.

The parameters that affect the observed drift in the baseline (i.e., asymptote and timescale, **Fig 2c,d**) remain rather independent of the experimental condition. This feature is more apparent in the ORIG dataset, but it still seems to hold in the FREQ dataset. An exception arises at the lower frequency (1 cpb) tested in the FREQ dataset. However, the case of frequency one is rather special and should possibly be considered as transitional between periodic and non-periodic stimuli.

**Fig 3** provides an idea of the quality of the fits by showing the *evidence* of the data in favor of the models tested (cf. [11,13,28]). Upper and lower rows correspond to Two-way and Global adaptation type respectively. For dataset ORIG, **Fig 3a** shows the logs of the odds ratio of the parametric model of **Equation 8** against a noise only model consisting of the block mean with variance similar to that of the data. Each bar is split into the log of the odds ratio of the full model to a drift only model that lacks the sinusoidal component (darker tone of the bars) added to the log of the odds ratio of the drift only model to the noise only model described above (lighter tone of the bars). This separation is possible because the models are nested so that the simpler models can be obtained from the full model by eliminating parameters. The evidence then compares the density of models likely to fit the data. **Fig 3b** shows the estimation of the frequency of the oculomotor response against the actual frequency of the stimulus for the three frequency values tested in dataset ORIG (3, 4, and 6 cpb). **Fig 3c** and **3d** shows the evidence and the agreement of the response with the five frequencies used in dataset FREQ (1, 3, 6, 12, and 24 cpb).

### State-equation fittings and model selection

To assess the generative model, we fit **Equation 5** to all data available. For illustration puposes only, we first show that the model provides a reasonable overall fit to the group data. **Fig 4** shows fits of the oculomotor response predicted by the full form of the generative model given by **Equation 5** with all five parameters described in the **Methods** section: *K, A, m, D* as well as the initial condition *G*. As before the qualitative agreement of the fits and the data is evident in both datasets. As we did with the phenomenological fits, we included the pre-adaptation blocks in each condition in each dataset.

**Fig 4.**
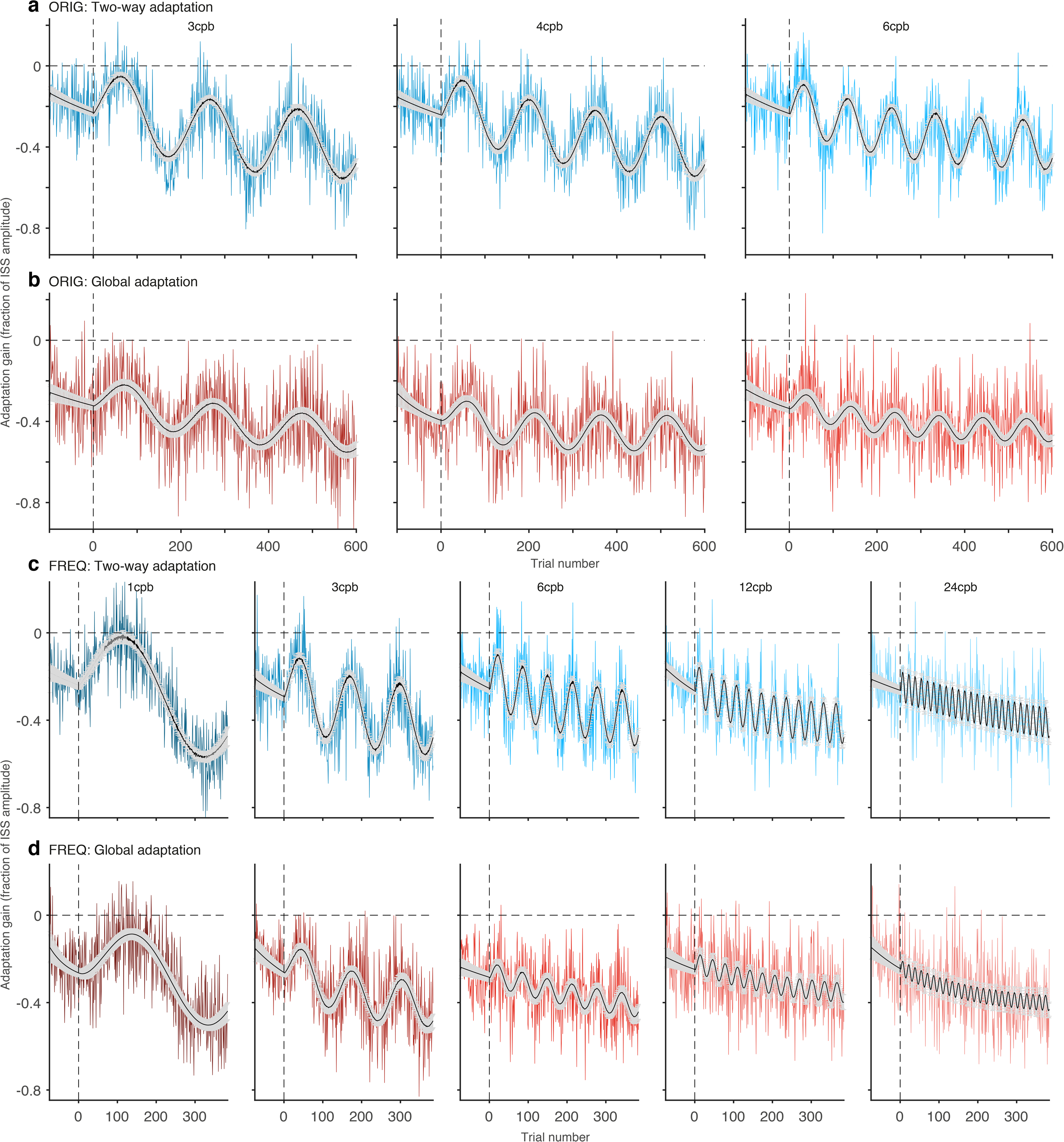
Fits of the oculomotor response predicted by the state-equation (Equation 5) with all five parameters. The plots show adaptation gain (colored lines), averaged over individuals in the (**a**) Two-way adaptation and (**b**) Global adaptation condition of the ORIG data set (reported in [11]), as well as the (**c**) Two-way adaptation and (**d**) Global adaptation condition of the FREQ data set, using the same paradigm over an extended range of frequencies. The stimuli input in the model fits is ***s***(***n***) (cf. **Equations 1–3**), which is zero in the preadaptation block. The same equation was fitted to data from each participant in each condition and experiment, to estimate parameters of the generative model on an individual bases. For illustration purposes only, the figure depicts fittings done over the averages along with 95% confidence intervals (gray shaded areas).

For all subsequent analyses, we fitted models to individual data. In particular, we compared 16 different models that differed from each other depending on which parameters were fitted (see **Methods** for details). We used Akaike’s information criterion (AIC) to explore statistical selection among these models. Akaike weights (cf. section II of [40]) are shown in **Fig 5** segregated by model and condition, for datasets ORIG and FREQ, respectively. In each condition (identified by adaptation type and stimulus frequency), we computed a matrix of weights in the following way. Because the best fitted model may differ between individuals, we first computed the AIC weights among the 16 models for each participant and condition. Then we averaged the resulting individual weights across participants. Results from this procedure are shown in **Fig 5**.

**Fig 5.**
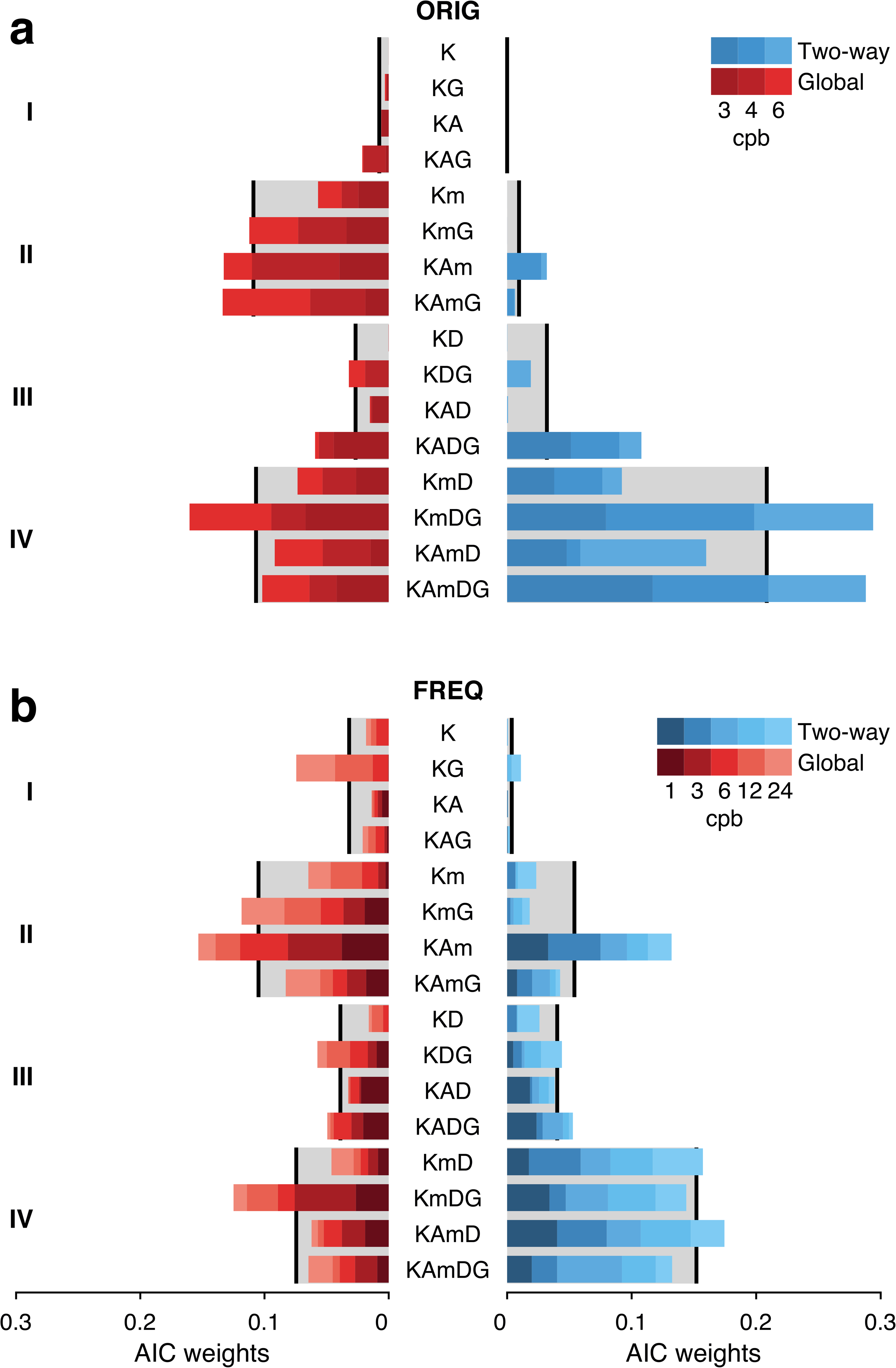
Akaike weights [40] for the 16 versions of the generative model, segregated by condition (frequency and type of adaptation). The label along the middle y-axis indicates the model for the weight displayed in the horizontal bars. Results from dataset ORIG (**a**) and FREQ (**b**). Weights for each of the three frequencies for each type of adaptation (blue tones for Two-way expanding to the right, red tones for Global to the left) are stacked for each model and color-coded as in **Fig 1**. The models are grouped according to the criterion described in the subsection *Rationale for generative model building and parameter exploration* in the **Discussion** section (see text for further details). Gray areas in the background indicate the average weight of the corresponding model group.

Inspection of **Fig 5** suggests clear overall preference for models in groups II (which include *m* but not *D*) and IV (featuring both *m* and *D*). We discuss below why this is expected on theoretical grounds given the features of the data. Models from group IV that learn based on two error samples, are preferred in Two-way adaptation, specifically the full model (*KAmDG*) and the model in which *A* was set to unity (*KmDG*). Models in group II that feature a single learning rate (error-correcting based only on the last experienced feedback), specifically *KAm* and *KAmG*, have an edge in Global adaptation. In what follows, we will focus on a comparison of these four models.

**Fig 6** shows the values of the generative parameters (Mean ± SEM, N = 10 for dataset ORIG, N = 13 for dataset FREQ) of the best models that learn only from the last experienced feedback error (*KAm,KAmG*). Upper and lower rows correspond to datasets ORIG and FREQ respectively. Learning rate *K*, persistence rate *A* and drift parameter *m* are shown in columns **a, b** and **c** of **Fig 6** respectively. **Fig 7** reports the parameters of the best models that update their hidden variable based on double error sampling. Those models are *KmDG* and *KAmDG*. **Fig 7 a-d** show respectively the learning rates *K* and *D* that weight the contributions of last and next-to-last feedback, the persistence rate *A* and the drift parameter *m*. Note that all models include the drift parameter *m* as a fitting parameter. We shall explain below why this should be expected.

**Fig 6.**
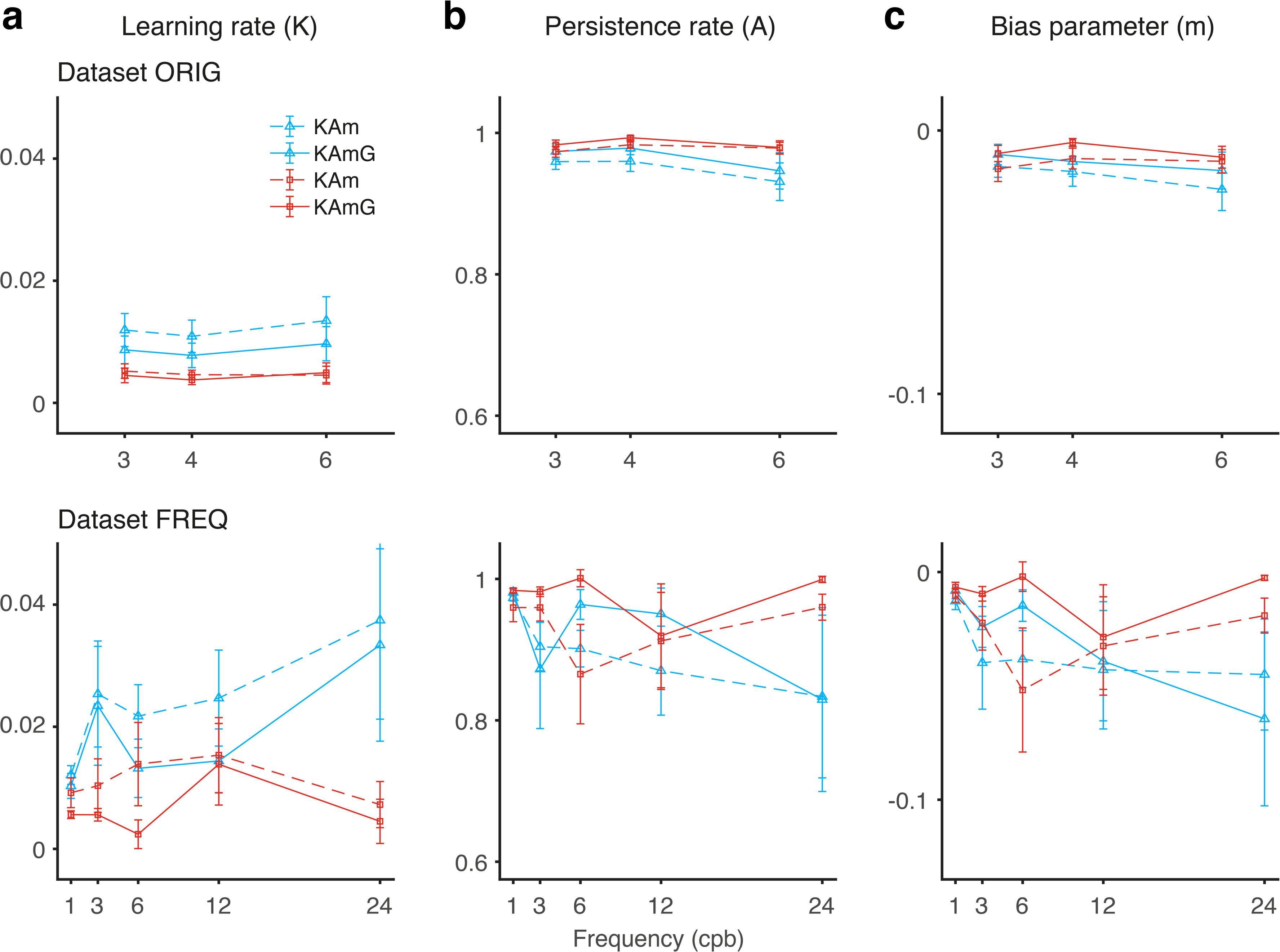
Average over individual parameters of generative models *KAm.KAmG*, the best among those that learn from the last feedback only (cf. Equation 5 with *D* = 0). Blue and red colors correspond to horizontal Two-way and Global adaptation, respectively. (**a**) Learning rate ***K***, (**b**) Persistence rate ***A***, and (**c**) bias or drift-parameter *m* are plotted as a function of condition. Both favored models feature ***A*** and ***m*** as fitting parameters. Note the variability in their fitted values across conditions, in particular for dataset FREQ. Error bars are ±SEM.

**Fig 7.**
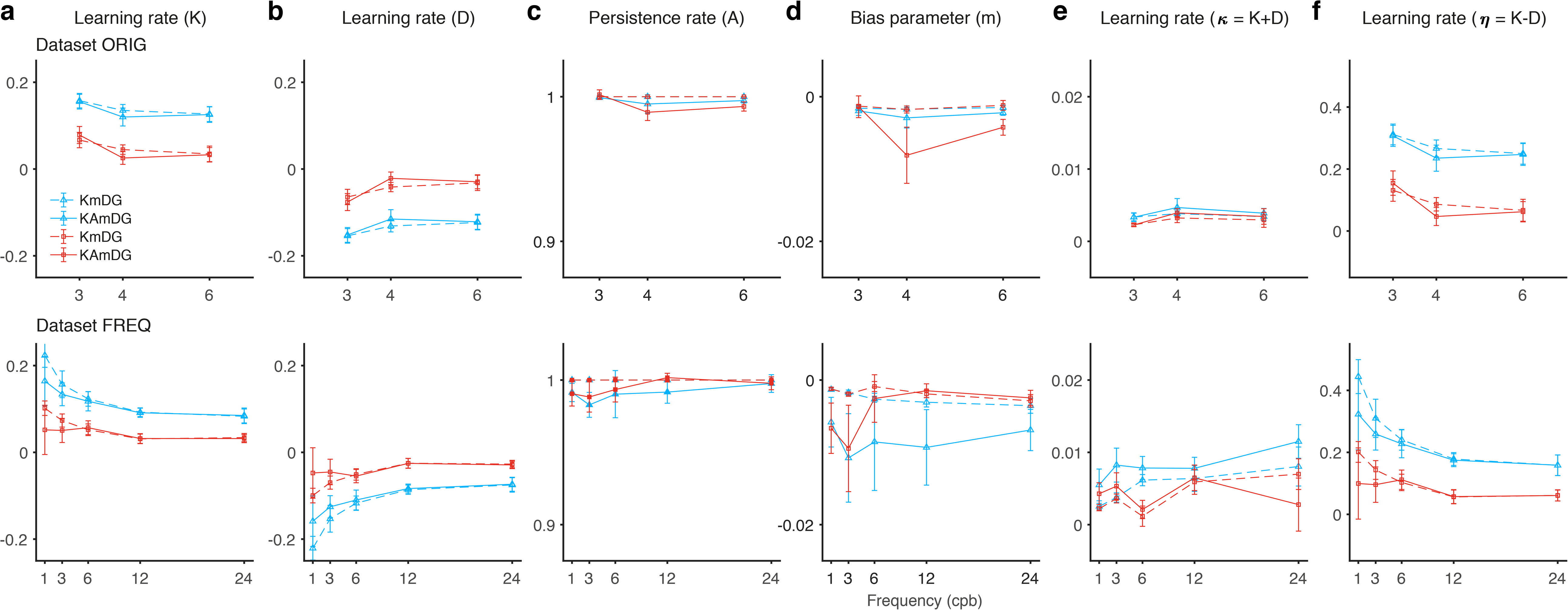
Average over individual parameters of generative models *KmDG, KAmDG*, the best among those learning from last and next-to-last feedbacks (double-error-sampling model; cf. full Equation 5). Blue and red colors correspond to horizontal Two-way and Global adaptation, respectively. (**a**) Learning rate ***K***, (**b**) Learning rate ***D***, (**c**) Persistence rate ***A***, and (**d**) bias or drift-parameter are plotted as a function of condition. Both favored models feature ***D*** and ***m*** as fitting parameters. Note that in both models, the bias parameter ***m***, and the persistence rate ***A*** in model ***KAmDG***, display much lower variability in their fitted values across conditions when compared to that of **Fig 6**. (**e,f**) Addition and difference of both learning rates, ***k*** = ***K*** + ***D*** and ***η*** = ***K*** + ***D*** (see text for discussion).

Again, several features are readily apparent from these plots. The learning rates (*K* and *D*) obtained from ORIG [11], show a rather clear segregation between Two-way adaptation and Global adaptation: *K* and *D* are larger for Two-way (blue colors) than for the Global case (red colors) suggesting that the extra variability brought upon by the random directions of the subsequent saccades characteristic of Global adaptation has a detrimental effect on all learning rates. They do not show a strong dependence on the frequency but the range of values used in that experiment was rather narrow, ranging from 3 to 6 cpb. This segregation in the learning rates between Two-way and Global adaptation is also clearly present in the best models fitted to dataset FREQ.

A feature observed in all cases is that in models that learn only from the last experienced error, the (single) learning rate (*K*) shows a mild increase with the frequency. This changes substantially if learning from the next-to-last feedback is included. In all of these models, the following features are observed. First, the magnitude of *K*, the learning rate of the last-feedback error-correction term increases by about an order of magnitude with respect to the models that do not have next-to-last error-correction. Second, the magnitude of the next-to-last error learning rate (*D*) is similar to that of the last error (*K*) but with opposite sign. This seems to suggest that the next-to-last error is weighted negatively (or actively attempted to be forgotten) in the algorithm. Third, the discrepancy in magnitude between *K* and *D* is consistently larger for Two-way than for Global adaptation (compare the separation between corresponding blue and red lines in **Fig 7, a** and **b**). Fourth, the learning rate *K* reverses its dependence with the frequency of the stimulus with respect to the models without *D*, and now decreases monotonically as the frequency of the stimulus increases. At the same time, the magnitude of *D* also decreases with the frequency. As a consequence, the discrepancy in magnitude between *K* and *D* is such that the addition of both learning rates approximately matches the range of the values of *K* fitted in the models that learn only from the last feedback (compare the values plotted in column **a** of **Fig 6** to those of column **e** in **Fig 7**). This suggests that when the additional error learning is not part of the model, the only learning rate fitted may represent an average across sub processes.

The values of the parameters fitted with the best four models are shown in **S1 Table** (Mean ± SEM, N = 10 for ORIG, N = 13 for FREQ). To assess dependence of the generative parameters on the experimental conditions we run 2 × 3 (ORIG) and 2 × 5 (FREQ) repeated-measures ANOVA on the fitted values using as regressors type of adaptation (Two-way and Global) and ISS frequency. Results are shown in **S2 Table** for the parameters given in **S1 Table**. We regard as more representative the results from dataset FREQ due to the more extended range of frequencies tested. Consistent with the qualitative observations mentioned above, while type of adaptation is highly significant for the learning rates in every model, frequency show significance for and only in the models that feature double error sampling (*KmDG*, for both datasets, *KAmDG* only for FREQ) but not in those learning just from the last feedback (*KAm,KAmG*). As for the persistence rate, frequency is never significant suggesting that it can be kept fixed as in model *KmDG*. Type of adaptation is significant in *KAm* and *KAmG* but such significance disappears in *KAmDG*.

### Analytical solution of the generative model: predicting the phenomenological parameters

The iteration of state-equations that learn from the last feedback already qualitatively predicts both components of the phenomenological response. In general, the complete response can be interpreted as a convolution of the stimulus with a *response function*. This response function integrates the stimulus by weighting the disturbance over a temporal window, the size of which depends on the magnitude of the learning and persistence rates that combine to assemble the weights (cf. **S1 Appendix**). Contributions from constant components of the disturbance that arise either from constant features in the stimulus (as in the traditional fixed-ISS paradigm [1]) or from intrinsic biases that may not be strictly error-based in nature (e.g., in our case represented by the drift parameter; cf. [37]) accumulate across trials, changing saccade gain in a monotonic fashion akin to a drift of the baseline towards an asymptote. Iteration of the systematically varying part of the disturbance results in its convolution with similar weights but the trial-by-trial variation usually prevents finding a closed form for the series re-summation. However, a sinusoidal disturbance avails a closed analytical integral solution, it is periodic with the same frequency, lagging the stimulus by a number of trials. Two new phenomenological parameters of this periodic response— its amplitude and lag—bare characteristic dependences on the learning parameters.

Above, we fitted the extended version of **Equation 8** to the data and obtained and reported estimates for its phenomenological parameters (i.e., frequency *v*, amplitude *a*, lag *ϕ*, asymptote *B*_0_, timescale *λ* and decay amplitude *B*; cf., **Fig 2** above). Similarly, we fitted the generative parameters for all generative models using the corresponding versions of **Equation 5. Figs 6** and **7** display those estimates for the four models that provided the best fits (excluding *D*: *KAm* and *KAmG* and including *D: KAmDG* and *KmDG* respectively).

When the learning algorithm includes several error-based terms, **Equation 5** can be integrated using techniques standard within the theory of LTIS [42]. This integration provides analytical predictions of the phenomenological parameters as functions of the learning parameters fitted with the generative models (**Equations 9** through **14**). We attempt matching these predictions to the values fitted using the phenomenological parameter estimation implemented before (see **Fig 2**). It should be pointed out, however, that the phenomenological parameter values have also been obtained from fits to the data and therefore should only be regarded as indicative reference values to guide intuition, not as ground truth. Validation of the actual underlying structure of the learning model relies ultimately on statistical model selection. Yet, a direct comparison between the fitted phenomenological parameters and analytical predictions evaluated on the fitted generative parameters is informative because a given value of a phenomenological parameter has to be compared to diverse combinations of the generative parameters depending on the specific structure of the learning model.

We start with **Equation 13** that provides a relationship between the expected asymptote of the adaptation gain at large trial number and the generative model parameters.

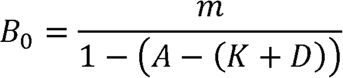

A first significant observation about this expression is that in order to observe a drift in the baseline of the adaptation gain (i.e., in order to have an asymptote *B*_0_ ≠ 0), a finite value of the drift parameter *m* is strictly necessary. If *m* vanishes, the adaptation gain would maintain a baseline pinned at zero regardless of the values of *K, A* or *D*. In addition, in a situation where *A*~1, 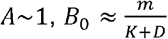 or 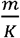 in models where *D* = 0. Note that these are all signed magnitudes, not absolute values. In other words, a small learning rate *K* or a small number resulting from the addition *K + D* will modulate the size of the asymptote and will determine its sign (i.e., will modify the degree of hypometria or hypermetria). Still a finite value for *m* is strictly needed to have non-zero asymptote. Recall that when *m* is not a fitting parameter, its value is set to zero. Due to the pervasive baseline drift across all of our data, all models favored under statistical model selection contain *m* as a fitting parameter. This is why model groups II and IV (cf. **Fig. 5**) are preferred, as pointed out above and in the **Discussion**. Note, in addition, that the smaller the learning rate (*K* or *K + D*), the larger the size of the asymptote *B*_0_.

Experimentally, we observed drifts towards higher hypometria in all averages and in most of the individual data. Note that formatting the data in terms of *adaptation gain* instead of *saccade gain* allows us to remove confounds coming from constant contributions from the stimulus and therefore the parameter *m* should be regarded as intrinsic to the system. In other words, *m* characterizes or quantifies learning that would occur in absence of stimulus disturbance (i.e., with zero ISS), as if the system has an intrinsic propensity to modify its gain by virtue of environmental or experimental conditions not necessarily linked to an error.

**Fig 8a** displays the matching of the analytical predictions of the asymptotes computed by inserting the fitted values of *m, A, K* and *D* into **Equation 13** for each participant’s data, to the phenomenological estimation of *B*_0_ obtained from **Equation 8** and the parameter estimation of the phenomenological fits of the data for both datasets and both adaptation types.

**Fig 8.**
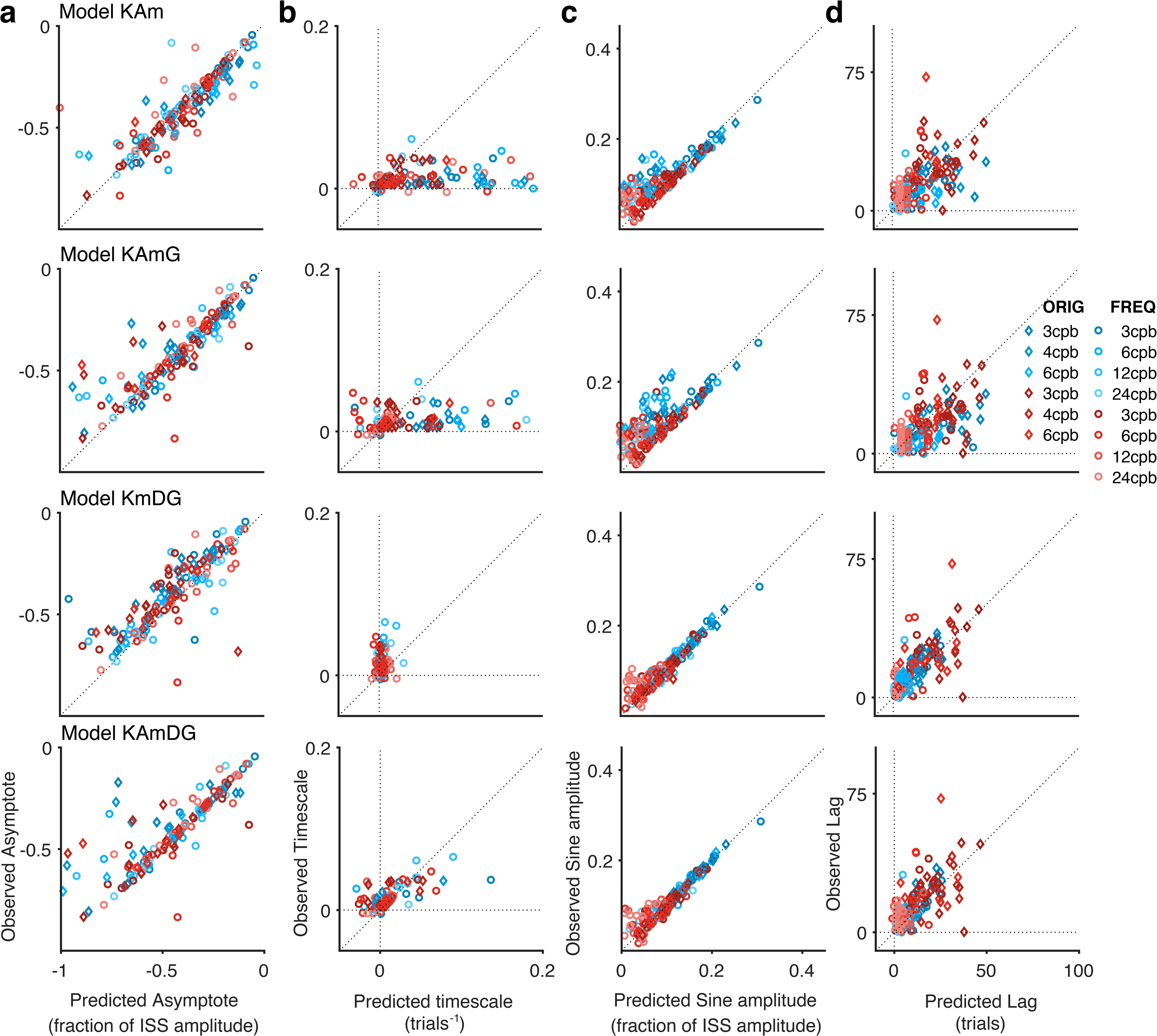
Comparison of phenomenologically fitted parameters to their theoretical predictions based on the generative model. y-axes show the values obtained with the phenomenological parameter estimation (**Equation 8**); x-axes show predictions obtained by inserting the best estimated values of the generative parameters into the the analytical expressions of **Equation 9–14** (cf. ‘*Using the generative model to predict the parameters of the phenomenological description of the adaptation gain*’ in the **Methods** section). Each row corresponds to one of the four best generative models. Each point is a single participant in a given condition and experiment. Data from the condition with a frequency of 1 cpb has been omitted because the predictions were poor for all models (in particular for the lag valiable, see the text for a discussion of this point). (**a**) Asymptotes of the gain at large trial numbers (***B_0_*** in **Equation 8** vs predictions by **Equation 13**). (**b**) Timescale of the decay of the baseline (***λ*** in **Equation 8** vs predictions by **Equation 14**). Note the wide spread of the predicted values for models ***KAm***, and ***KAmG*** that also results in several ouliers beyond the limits of the subplot. In contrast, models ***KmDG*** tend to underestimate the phenomenological timescales. Model ***KAmDG*** provides the best prediction of the phenomenologically fitted values with only two points just beyond the limits of the plot. (**c**) Amplitude of the periodic component of the gain (***a*** in **Equation 8** vs predictions by **Equations 10–12**). Note that models ***KAm*** and ***KAmG*** tend to underestimate the observed amplitudes of the peridic component of the gain. (**d**) Lag of the periodic component of the gain (*ϕ* in **Equation 8** vs predictions by **Equations 9–12**). The plots reveal a slight tendency for models ***KAm*** and ***KAmG*** to overestimate the length of the lag with respect to the predictions of the models including double error sampling (***KmDG*** and ***KAmDG***).

A second parameter characteristic of the baseline drift is given by the timescale. **Fig 8b** shows predicted values for the timescales that result when values of *m, A, K* and *D* fitted with the state-equation are inserted in **Equation 14**. The first two rows of **Fig 8b** show a clear overestimation of the baseline timescale in models that do not feature double error sampling (i.e., *KAm* and *KAmG*) as several individual data points fall outside the boundaries of the plot. Yet, models that include corrections based on the next-to-last error term, seem to underestimate the timescale (in particular model *KmDG*). When introducing **Equation 14**, we pointed out that if the second error learning rate *D* is negative, the dominant mode in the solution still features a monotonic decay that can fit the phenomenologically observed exponential baseline drift of the gain. This is indeed the case in the majority of fits to the individual participants’ runs: Across models in group IV, *D* was non-negative in only 13% of the individual runs; 6% for Two-way adaptation and 21% for Global adaptation data. For model *KmDG, D* was non-negative in 7% of all runs; with only 1% (1 run out of 95) for Two-way adaptation and 14% for Global adaptation. Furthermore, when estimating the timescales of models that include double error-correction, **Equation 14** consistently gives smaller values than for models without the second error term (cf. compare subplots of **Fig 8b** for the corresponding models, and **Fig S1** in **S1 Appendix**). This ordering relation between the timescales of models with and without *D* was unknown before conducting the fits. Thus, data collected using a sinusoidal adaptation paradigm suggests that including a second error-correction term yields a significant decrease in the timescale with respect to models featuring a single error-correction term. Therefore, the integration window (i.e., the inverse of the timescale) of models with double error-correction grow significantly larger compared to those that lack the second error sampling.

Asymptote and timescale are parameters traditionally investigated and reported in adaptation to fixed-step disturbances. Sinusoidal adaptation paradigms provide two additional parameters associated to the periodic component of the adaptation gain observed in these protocols. **Fig 8c** and **d** compare predictions for the amplitude and the lag of the periodic component of the gain obtained by using **Equation 9** through **12** above. Data from both datasets suggest that models that do not feature double error sampling underestimate the magnitude of the amplitude of the periodic component of the oculomotor response (cf. predictions from these models in **Fig 8c**). This feature in fact is common to all models that learn from a single feedback and include *m* (besides models *KAm* and *KAmG*; not shown) but the inclusion of *D* helps mitigating misestimation of this amplitude.

The last comparison is provided by the lag of the periodic component. **Fig 8d** compares predictions based on the state-equation learner (**Equations 9** and **11** furnish predictions for the components of the lag *ϕ* and *φ* after inserting the parameter values fitted with **Equation 5**) and the phenomenology (parameter *ϕ* in **Equation 8**). From **Fig 8c** and **d** it is apparent that the models that include both and as fitting parameters provide better predictions, also displaying less variability across participants, in particular for the Two-way adaptation type. Among models with *D* = 0, again models *KAm* and *KAmG* fit best. **Fig 8d** shows, however, that these models appear to overestimate the lag (cf. compare corresponding subplots in the figure), while models that have a second learning rate *D* match better the empirically observed lag. In addition, all models fail the estimation of the lag for a disturbance of frequency one as they all significantly overestimate the lag observed. Even though the predictions of the other phenomenological parameters are reasonable (the amplitude of the periodic component, timescale and asymptote), predictions for the 1cpb condition for both Two-way and Global adaptation have been omitted altogether in **Fig 8**. This mismatch between the direct phenomenological estimation of the lag from the data and the analytical predictions arising from the integration of the state-equation for the case of the 1cpb condition, may be rooted in the fact that the functional dependence of the phenomenological parameters on the generative ones is determined by the specific sinusoidal dynamics of the driving stimulus, while the case of a 1-cpb frequency is the least periodic condition among all tested.

## Discussion

We used a modified version of the traditional two-step saccade adaptation paradigm ([1]; see [2,46] for reviews) in which the size of the second step varied as a sinusoidal function of trial number with an amplitude of 25% of a fixed pre-saccadic target amplitude. We recorded observers’ eye movements at a total of six different frequencies and applied the sinusoidal disturbance always along the saccade vector which was aligned either in a horizontal bi-directional fashion (Two-way adaptation) or in random directions drawn from a uniform circular distribution (Global adaptation). The oculomotor response, quantified by the adaptation gain, followed the disturbance variation with comparable frequency, an amplitude ranging between 10 and 30% of that of the stimulus (i.e., 2.5 to 7.5% of the saccade amplitude), and lagging the stimulus by a few trials. In addition, it developed a systematic drift of the baseline towards larger hypometria that reached asymptotes of around 40% of the disturbance amplitude (i.e., 10% of the saccade amplitude) and was largely comparable across conditions. The phenomenological description in **Equation 8**—composed of a periodic response and an exponential decay—captured this behavior well and we estimated all six parameters pertaining to that description.

The present study explored whether the phenomenology described by **Equation 8** can be modeled with a state-equation, i.e., a generative rather than descriptive model of the underlying sensorimotor learning. We clearly show that the recorded saccade adaptation data is indeed predictable in a robust and stable way using a linear time invariant state-equation similar but not identical to those proposed before in the literature. Moreover, in previous accounts, simulations based on generative models as well as ad-hoc fittings (mostly exponential or monotonic) of the temporal evolution of the gain were provided without specifying a pathway of how to evolve from one description to the other. We suggest that connection here and provide results of the derivation involved in transitioning between these descriptions.

### Rationale for generative model building and parameter exploration

In mathematical terms, the functional form in **Equation 8** is the integral solution of a family of LTISs of which **Equation 5** is a particular example. It is referred to as a state-equation or state-space model because the internal variable *x* characterizes the gain or state of adaptation of the system. This algorithm is *generative* because it estimates the value of *x* at trial *n* + 1 by modifying its estimate at the previous trial including possible effects of systematic biases and correcting the former value by weighting sensory feedback resulting from movement execution [21,26,47,48] (see also [25,32]; for further details on our specific use see the **Methods** section). Here we limit our discussion to *noise-free* generative models in that **Equation 5** does not include any noise term. Yet, **Fig 1** together with **Fig 4** suggest that integral solutions as well as numerical outcomes of noise-free generative models survive ensuing variability, at least for the paradigm, type of stimulus and within the ranges of the conditions tested.

We analyzed 16 models that differed in the specific parameters that were fitted and then used Akaike’s information criteria to attempt model selection. Since we were primarily modeling intrinsic error-based sensorimotor learning, the learning rate *K*—that weights the impact of the last feedback error on the state of adaptation—was present in every model. Second, we included the initial condition *G* as a fitting parameter in half of the models. This parameter is not part of the trial-by-trial learning algorithm and its effect should decay as the trial number increases (cf., **Equation 7**). However, the initial condition affects the amplitude of decay of the baseline drift (cf. *B* in **Equation 8**). Because the argument of **Equation 5** is an internal variable not directly experimentally accessible, a proxy for its initial value can only be approximated (for example, by averaging the first five gain values in the block) or included as a fitting parameter. Third, we included a persistence rate *A* that weighted how much of the estimate from the previous movement remained in the subsequent one. The fourth parameter, *m*, captured systematic effects, that are not error-based in nature, and gave origin to drifts in the baseline that were pervasive across all conditions. Finally, we considered the plausibility and study the effects of a second learning rate *D* that tracks errors other than the most recent (here, the next-to-last feedback error).

To further discuss the effect of the generative parameters, we split the 16 models into four groups:

I. Models that neither included terms depending on the second learning rate *D* nor the drift term *m*(*K, KG, KA, KAG*);
II. Models without terms depending on *D* but including *m*(*Km, KmG, KAm, KAmG*);
III. Models including terms depending *D* on but excluding *m* (*KD, KDG, KAD, KADG*);
IV. Models with both *D* and *m* terms included (*KmD, KAmD, KmDG, KAmDG*).

We recall that in models where *A* is not a fitting parameter, *A* = 1. The groups are listed on the left side of **Fig 5**. Models within group I consistently fitted worst. Moreover, models that do not include *m* (groups I and III) cannot capture an evolution of the gain into a stationary asymptotic value because the state equation does not admit a solution featuring that behavior (that is, if the stimulus has no constant term). These models, however, may be useful in experimental paradigms where a stable state of adaptation is not clearly reached either because the length of the adaptation block used may be too short or because the driving disturbance is unbounded (for example a linear ramp). On the other hand, models that include sampling from two errors (cf. groups III and IV) will likely be better suited to extract correlations built into the stimulus as it is the case of a sinusoidal ISS.

The fits of the phenomenological model (**Fig 1; Equation 8**) suggest that asymptotic behavior of the baseline and reflection of the stimulus self-correlation (entraining) were clear structural properties of the oculomotor response. The analytical solutions of models in both groups II and IV are consistent with this phenomenology. **Fig 5** summarizes the AIC weights emerging from the fits to the individual participants’ data. The weights shown in the horizontal bars are averages over individual participants’ weights for each condition and color coded by the frequency of the stimulus. Data from Two-way adaptation is depicted with blue tones in bars increasing towards the right. Global adaptation is shown with bars spanning to the left in red tones. The average weight for each model family is shown by the gray background behind the corresponding group. While models in group II already generate responses in qualitatively good agreement with the evolution of the adaptation gain, it remains to be decided whether corrections based on the memory of more than a single error provide for a better fit. AIC weights show that group IV clearly outperforms all others in Two-way adaptation in both datasets, suggesting that the best generative model to describe this type of adaptation includes all four parameters *K, A, m* and *D*. In Global adaptation, models from group II either match or slightly outperform those of group IV. Model comparison showed that a state-equation including a single parameter or any combination of only two of the four parameters *K, A, m* and *D* could not adequately account for our data (cf. **Fig 5**). In addition, an inspection of actual values of the parameters fitted across the population suggests that the parameters and may be set to constant values, that is, to almost one for the former and to a very small and negative number for the latter (cf. **Fig 7**, columns **c** and **d**), at least within the range of frequencies tested in these experiments. Overall, the drift parameter *m* and the second learning rate *D* proved useful and necessary to account for systematic effects in our data, suggesting (1) that some changes in the adaptation state are not error-based and (2) that—at least under specific circumstances—the brain keeps track of at least one extra occurrence of the error besides the last experienced one. Three-parameter models that did not involve *D* (specifically *KAm*) were most successful in Global adaptation and in the high frequencies of Two-way adaptation. This could be simply a reflection of increased levels of measurement noise in these conditions giving an upper hand to models with fewer parameters. More interestingly, it could point to an architecture that samples two errors only under certain conditions, for instance, when errors are repeatedly experienced for the same saccadic vector, or, when the variation of the feedback error has a high signal-to-noise ratio. We speculate that overtraining along a given direction, understood as the repetitive experience of consistent error along similar saccade vectors in Two-way adaptation (note that in our paradigm Two-way adaptation stimulates only two retinal locations) may give rise to vector specificity and, consequently, to the adaptation fields typically observed with fixed-step paradigms. Indeed, Rolfs and collaborators [18] suggested that Global adaptation, featuring apparent full transfer across random directions, appears to onset ahead of the development of vector-specific adaptation fields. This appears consistent with the present finding that models that rely on a single error-correction show timescales corresponding to faster evolution of the baseline drift (although with longer lags in the sinusoidal component) as compared to those of Two-way adaptation (featuring shorter lags in the sinusoidal component consistent with tracking the stimulus more closely due to the repetitive training in a specific direction).

### Drift in the baseline and the meaning of m

The persistent drift of the baseline towards higher hypometria is a distinctive feature in our data that cannot be accounted for on the basis of motor adaptation [49]. We included an extra parameter *m* to account for this drift in mean adaptation gain towards an asymptote differing from the mean of the stimulus (cf. **Equation 13**). This parameter is conceptually novel, distinct and independent of the persistence rate *A*, and determines the presence of a non-zero asymptote via **Equation 13**. Because in our paradigm the goal of the task was to land on the target as close as possible, and because the sinusoidal ISS introduced a continuously changing prediction error, the best expected outcome would be to track the disturbance within the levels of error typical of trials without disturbance. With respect to that goal, the presence of a baseline drift introduces an additional discrepancy that does not, however, hinder successful adaptation to the disturbance.

Saccadic eye movements slightly undershoot their target on average [50] and this systematic offset corresponds to the internally predicted visual outcome of a saccadic eye movement [51,52]. We surmise that our paradigm may have yielded a re-calibration of this desired offset [53] over the course of an experimental run. This recalibration towards a larger undershoot may result from the high probability of a quick return saccade after every eye movement in our fast-pace paradigm, reducing the utility of maintaining a saccade gain close to one. We note that this systematic decrease in saccade gain may in general—albeit to different degrees—pervade the study of saccadic adaptation (but see [7,54]). In fixed-step paradigms (as opposed to the sinusoidal paradigm employed here), however, it would have been obscured as the error-based correction for the surreptitious target displacement undergoes similar dynamics as the drift reported here.

On the other hand, from the point of view of the internal model of the movement that the brain may implement [33–35], this bias parameter *m* may hint to a discrepancy between the experimental coordinate system where measurements are acquired and the coordinate system in which the internal model is represented.

On a neurophysiological level, the small systematic bias that gives rise to the drift of the baseline may originate from the dynamics of the responses in the neuronal substrates involved with saccade adaptation ([55–60], Reza Shadmehr, *personal communication*, July 12, 2018). It is also possible that the fast-pacing used in our paradigm exacerbates effects that generate a small and negative bias parameter, *m*, which appeared to onset already at the pre-adaptation block. That would further suggests that the magnitude of *m* may depend on the inter-saccade interval as well as on the precise timing of the ISS onset, which should be addressed in future studies.

### Consequences of learning from double error sampling (D parameter): Two learners?

The models that best explained the data featured a double error sampling, learning not only from the feedback experienced after the last saccade but also from the movement that occurred in a trial before that. Hence, the best models used a feedback reaching further back in time through the *K*- and *D*-terms of **Equation 5**. Yet, does the oculomotor system actually implement this double error sampling that may coherently participate in a single internal model prediction? We suggest that the brain may attempt to approximate the performance achieved by the double-error-sampling algorithm by using two single-feedback learners operating on appropriate combinations of the stimulus sampled at two different times.

To understand that, we return to **Equation 5**. For simplicity, we will assume that *m* = 0.

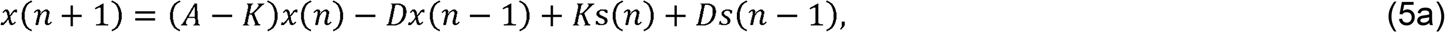

and write a transformation among state variables sampled at two different trials as,

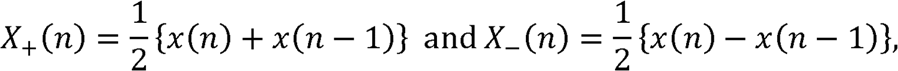

that can be substituted in the RHS of **Equation 5a** using the inverse relations:

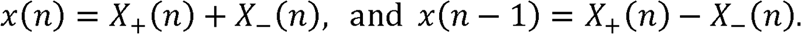

We can re-write **Equation 5a** in terms of these alternative state variables *X*_+_ and *X*_−_:

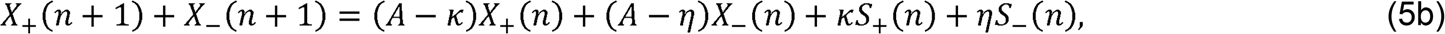

where we adopted the definitions of 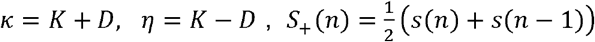 and 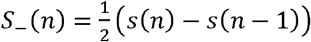. **Equation 5b** avails the interpretation of the generative model as selectively learning into two component channels that learn from a single feedback error taken from different sources. The source for the learner *X*_+_ is the mean of the two samplings of the stimulus, i.e., 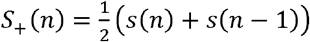. The source for the second learner is the rate of change of the stimulus across the sampling events given by 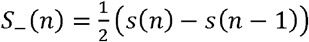 which, when the samplings occur on successive trials, it could be interpreted as the discrete time derivative of the stimulus taking the elementary timestep as the (average) inter-trial interval.

Note that the representation in terms of these alternative internal variables would significantly alter the underlying structure of the noise-free learning model. But if we insist on keeping a close connection to the parameters extracted using the double-error-sampling algorithm, we would expect that the learning rate for learner *X*_+_ would be the addition of the rates for the two errors, *k* = *K + D*, while for learner *X*_−_ it would be *η* = *K – D* (cf. **Fig 7**, columns **e** and **f**). In all our fittings using the double error sampling, *K* and *D* were very close in magnitude but carried opposite sign. Furthermore, *k* was small and similar in magnitude to the learning rate *K* obtained for models that learned only from the last error. Because *D* was negative, the learning rate *η* for the second learner became also positive but much larger than *k*, in fact about an order of magnitude larger (**Fig 7**, columns **e** and **f**) effectively enhancing the overall gain of the process without driving the system unstable [61–63]. As a consequence, (*A – k*), which can be thought of as an *effective A*_+_ will be much closer to unity than *A*_−_ = (*A – η*). Therefore, *X*_+_ will learn and forget much slower than *X*_−_.

Using this double error sampling, the oculomotor system could track the rate of change of the stimulus from one saccade to the next, besides just its last change in size and it would approximate the learning efficiency of the double-error-sampling algorithm. The new internal learning variables (*X*_+_ and *X*_−_) would learn from smoothed-out versions of the disturbance resulting from the average sum and difference of the two sampled inputs. Whether this constitutes an advantage over learning exclusively from the last feedback depends on the nature of stimulus. If the disturbance is constant or fully random there would be very little advantage in performing the double error sampling. In the former case, the inter-sampling variation is zero leaving nothing to learn. In the latter, the inter-sampling variation would be another random magnitude and there would be little advantage in learning from the variation in the feedback. However, if the mean of the disturbance varies in a systematic way—as it does during sinusoidal adaptation, and presumably in natural scenarios—learning from its rate of variation would be advantageous and could well justify a large learning rate. In the representation of the double-error-sampling model, unlearning actively the next-to-last sampled feedback error (i.e., with a large and negative *D* subtracted from an enhanced *K*) would materialize this advantage with little extra investment. However, a negative learning rate feels counter-intuitive as learning is believed to follow the direction of the correction suggested by the feedback. Segregation of the learning underlying motor (or saccade) adaptation into two learners displaying similar characteristics to those suggested here have indeed been proposed in other contexts [8,25,64,65]. The argument presented above suggests a mechanistic way to construct a two-learner system, in which the components *X*_+_ and *X*_−_ can be considered statistics in counterphase. To approximate the double-error-sampling learner, the system may hold in memory both samples, compute mean sum and differences between the samples and implement two learners based on those statistics rather than from bare values of errors or stimulus occurrences. To achieve that, the oculomotor system would need to keep memory and weight prediction errors from a former time scale besides the last feedback [65].

An important point to notice is that, even if there is double error sampling, it does not need to be strictly the next-to-last error. It would be enough that the brain keeps a correlation of errors over two different trials (cf. [66]) although it would be reasonable that they are spaced only by a short delay [61]. This is a reasonable generalization since the inter-trial interval is rather arbitrarily set by the pacing of the task that may or may not match a possible internal sampling frequency by the brain. The frequency of the stimulus then determines to what degree differences in the stimulus can be sampled, which may explain the dependence of the amplitude and lag of the periodic component of the response with the frequency as well as the fact that the evidence for the full model seems to peak at intermediate frequencies. In other words, it may be easier to learn at certain frequencies (for a fixed amplitude) or at certain effective rates of change of the stimulus.

### Dependency of learning rate on perturbation dynamics: Linear but not strictly Time Invariant Systems

We further explored whether the values of the generative parameters exhibited dependence on the experimental condition, specifically with the type of adaptation and the frequency of the disturbance. The parameters of our models remained time-invariant across pre-adaptation and adaptation blocks. However, we did not rule out that these parameters may change with adaptation type and stimulus frequency. In fact, LTIS models with parameters not strictly time-invariant have been invoked to model (meta-learning in) savings in adaptation to visuomotor rotations [32]. Strict LTIS models with two learners had been able to successfully account for savings in long-term saccade adaptation [8,25,64,67]) but were not able to fit differences in the dynamics of the adaptation, extinction and re-adaptation phases observed using counter-adaptation and wash-out paradigms in adaptation to visuomotor rotations without letting the rates change across the phases [32].

We limit our discussion to the best four generative models selected in the **Results** section. In models *KAm* and *KAmG* (and in general in all models of groups I and II), the (only) learning rate *K* remained relatively independent of, or exhibited a tendency to grow with, the frequency of the stimulus (**Fig 6a**). Learning rates for Two-way adaptation roughly ranged between 0.01 and 0.035 fraction of the error across the frequencies tested. The same parameter in Global adaptation was smaller and remained within the range 0.005 to 0.015 (cf. **Fig 6a** and **S1 Table**). These observations were confirmed by ANOVAs run on the fitted values of the parameter *K* in models *KAm* and *KAmG* in that type of adaptation was always a significant factor while ISS frequency never was (**S2 Table**). These values of *K* compare reasonably well with the magnitude of learning rates previously reported in the literature (cf. [8,19]). The dependence of the learning rate on the frequency of the disturbance seems in qualitative agreement with results from reaching experiments in which subjects learned to track a target undergoing surreptitious displacement that followed a random walk [30,47]. Using a Kalman filter to estimate corrections to the learning rate due to various types of variability Burge and collaborators [30] argued that the learning rate increased as the drift of the walker increased. In the sinusoidal adaptation paradigm where the amplitude of the sine function that produces the ISS is of fixed amplitude, this situation occurs when the frequency increases because its size from one trial to the next changes faster. However, this suggestion seems at odds with the intuition that a more consistent stimulus should drive more efficient adaptation [68,69]. In particular, it has been reported that a smooth gradual variation results in more efficient adaptation [3,70]. If this were the case and reflected onto the model parameters, the learning rate should be higher for smaller frequencies.

However, the dependence of the learning rate(s) on the frequency described above changed rather dramatically when double error sampling was included (cf. **Fig 7**, columns **a** and **b**). Interestingly, in models that feature double error sampling, the learning rate of the most-recent error-term (*K*) reversed its tendency and decreased as the frequency increases, achieving its highest values in the conditions of lower frequency, this is, in situations where the stimulus displayed higher consistency. Concurrently, the learning rate for the next-to-last feedback (*D*) achieved its most negative values at lower frequencies and grew less negative as the stimulus frequency increased. In the alternative scenario of two additive learners with single error correcting terms that learned respectively from the half-sum and the half-difference of the two sampled errors suggested in the previous sub-section, the learning rates *k* and *η* also showed a distinct dependency on the ISS frequency. The *slow-learner* (with learning rate *k*) would only have corrected up to 1% of the average of the two errors sampled while the *fast-learner* (with rate *η*) would have produced corrections of up to 40% of the change experienced between the two sampled errors (cf. **Fig 7e** and **f**). Note that this massive change in the dependence of the learning rates on the frequency was a consequence of changing the hypothesized structure of the model and not of correcting the magnitude of the rates for effects of variability. Once again, ANOVAs confirmed that not only the type of adaptation but also the stimulus frequency had significant impact on the learning rates (*K* and *D*, as well as *k* and *η*) in models *KmDG* and *KAmDG* as well as all models of group IV (cf. **S2 Table**).

In contradistinction, the retention rate *A* (**Figs 6b** and **7c**) and the bias parameter *m* (**Figs 6c** and **7d**) remained relatively independent of the frequency under such changes, although their overall variability was clearly reduced in the models featuring double error sampling (contrast the value ranges of *m* and *A* in **Fig 6**, against the corresponding ones in **Fig 7**, aside from model in which *A* = 1; see also corresponding entries in **S1 Table**). ANOVAs run over these parameters further confirmed non-significance of the frequency except for model *KmDG* on *m* in dataset ORIG (**S2 Table**). Type of adaptation occasionally modulated *A* in dataset FREQ in models with a single error term. Taken altogether these suggests that both *A* and *m* may be largely frequency independent and can be modeled as constant values maybe differing in value for Two-way and Global types.

In summary, introducing a second error term increased the magnitude of both learning rates (*K* and *D*) by an order of magnitude with opposing signs. The learning rates of these models showed a clear dependence on the frequency of the disturbance: higher stimulus consistency (i.e., lower stimulus frequencies) correlated with higher adaptation efficiency. At the same time, the inclusion of the double error sampling reduced variability in the estimates of the persistence rate *A* and the drift parameter *m*, indicating that their estimates were not affected by the ISS frequency, and could thus be set to appropriate constant values.

### Relation to previous work on sensorimotor control and adaptation

Multiple distinct learning processes contribute to sensorimotor adaptation [8,25,64–66,71]. Recent research conducted primarily within adaptation to visuomotor rotations or in reaching movements, suggests that adaptation can be decomposed into two fundamental processes that may operate in parallel: one that would be responsible for implicit learning that progresses slowly and can be described mechanistically by a state-equation [49]. This slow learning process is relatively stable over breaks, takes place with automatic, reflex-like behavior and its properties tend to be sturdy and do not change fast with recent experience. A second, parallel process, in turn, learns explicitly, is faster although it may require longer reaction time and possibly voluntary behavior to be engaged. This faster process would exhibit longer term memory of prior learning [71–74].

We believe that our paradigm taps only the first, implicit component. Yet, we suggest that our analyses provide evidence for two separable subcomponents, although both would be intrinsic in nature [75]. In fact, a key difference between our oculomotor learning and learning that occurs in adaptation to visuomotor rotations and during reaching in force fields is that our participants were primarily unaware of the inducing disturbance. In contrast, in the aforementioned paradigms, participants immediately notice a disturbance even when they may not be fully aware of the exact effect. In this sense, our paradigm could be considered qualitatively closer to that used by Cheng and Sabes [22] who studied calibration of visually guided reaching in participants fully unaware of the stimulus manipulation. Their paradigm used a random, time-varying sequence of feedback shifts. They found that a linear dynamical system (LDS) with a single error term and trial-by-trial state update for variability implemented with an estimation-maximization algorithm successfully described mean reach point and the temporal correlation between reaching errors and visual shifts. They further argued that the learning taking place under a random stimulus generalizes to a situation of constant shifts in a block paradigm and, therefore, that adaptation dynamics does not rely on the sequence (or correlation) of feedback shifts but can be generally described with the LDS model. In contrast to random or block constant ISS, our paradigm featured a disturbance that was fully self-correlated since it followed a sine variation with the trial number. Therefore, it may prove advantageous for the oculomotor system to extract correlations embedded in the disturbance because they would help tracking the target. As pointed out, including double error sampling would serve this purpose.

We believe that the presence of a systematically varying disturbance enables a further decomposition of the implicit component of adaptation, perhaps into a primary one, that attempts to mitigate the end-point discrepancy regardless of self-correlations in the disturbance, and a second one that attempts to extract (and use) such correlations. It remains an open question how these putative subprocesses may map on distinct or overlapping anatomical structures, such as cerebellar cortices, deep cerebellar nuclei and extracerebellar structures [55,57,59,60,64,76–80].

A recent study suggested that learning in dynamic environments may be adequately modeled with an algorithm popular in industrial process control, the proportional-integral-derivative (PID) controller [81]. The algorithm generates a control signal adding three error-related contributions: a term proportional to the current error that resembles a usual delta-rule (the P-term), a term that integrates over a history of errors experienced before the current one, and a derivative term estimated from the difference between the last two errors. The model shares some features with ours, in particular that the learning rate for the next-to-last error needs to be negative to approximate the derivative term. The PID controller acts on the actual recorded errors (the equivalent of the visual errors observed after each saccade is executed) and contains no internal state estimation, whereas our model operates on an internal variable that contains the state estimation of the prediction error that would result from the movement execution. Our state variable in fact accumulates and retains a substantial portion of the history of previous error (the persistence term in **Equation 5**, see also the example given in **S1 Appendix**), which is updated on a trial-by-trial basis by the term proportional to the latest prediction error (the delta-rule term). The inclusion of an extra error in our state-equation (specifically that of the previous to last one) effectively brings into play a contribution similar to the derivative term of the PID model. In short, our *D*-term enables a systematic correction to the integral term (our *A*-term) that otherwise would be determined rigidly by the iteration of the equation. In that respect, keeping track of former errors enables a structural correction that acts at a global level even when it is introduced on a trial-by-trial basis, lending both robustness and flexibility to the algorithm. Ritz and collaborators [81] further compared the performance of the PID model to a Kalman filter used to update a state variable in presence of noise applied on the single error structure of the usual delta-rule and found that the PID controller performs better. A further similarity with the aforementioned work lies in their observation that models with a derivative term are usually not readily selected under statistical model selection even when they may display significant improvement in the description of the behavior (see [81] for a longer discussion on this point).

## Conclusions

Having adequate generative models that describe eye movements have been stressed before [80,82–86] as an important tool to assess, at the single patient level, a variety of movement abnormalities that have been identified as markers of neurological conditions or disorders at a group level. In this study, elaborating on the idea of tracking a memory of errors [65], we attempted to identify and constrain a relatively minimal set of requirements that would suffice to model saccade adaptation data collected under the paradigm and stimulus that we recently implemented [11] but that would also include previous accounts of the phenomenon under other known paradigms. While certainly many refinements are still due, we unveiled features of an algorithm that seems suitable to account for the sensorimotor learning observed in our data. We hope it can be generalized, extended and adapted for use in future research.

## Acknowledgements

We thank Thérèse Collins and members of the Rolfs lab for insightful discussions and help with data collection.

---

**S1 Table. Generative parameters fitted with the best four models.** Model name is shown at the top. The corresponding datasets can be identified by the stimulus frequencies tested: ORIG: 3, 4 and 6cpb. FREQ: 1, 3, 6, 12 and 24cpb.

**S2 Table. ANOVA results on the generative parameters fitted with the best four models.** Repeated-measures ANOVA (2 × 3 on data from ORIG; 2 × 5 on data from FREQ) with factors type of adaptation and stimulus frequency was run on each of the four best models. Model name is shown at the side of the table and parameter names are on the top. The dataset is indicated in the cell at the upper left corner next the the parameter names. Highlights indicate the cases where the corresponding factor shows significant effects.

**S1 Appendix. Predictions of a state-equation with a sigle, most-recent error-based correction term. Effects of including next-to-last error-sampling.**

