## Supplemental Appendix for "A generative learning model for saccade adaptation"

**S1 Appendix. Predictions of a state-equation with a single, most-recent error-based correction term. Effects of including next-to-last error-sampling.**

*Iteration of a $KAmG$ model*. We can obtain an idea of the expected responses by iteration of **Equation 5**. For simplicity, we drop the term involving the second-to-last error ($D = 0$) to obtain:

$$x(n+1) = \left[G(A-K)^n + \{1-(A-K)^n\}\frac{m}{1-(A-K)}\right] + K\sum_{i=1}^{n} v(i)\,(A-K)^{n-i}. \qquad (A1)$$

We have inserted **Equation 3** so that $v(i)$ represents the portion of $s(n)$ that depends on the trial number. For sinusoidal adaptation datasets, $v(i) = p(i) \cdot \sin\left(\frac{2\pi f}{N} i\right)$. **Equation *A1*** can be split into two major contributions, one involving only constant features of the disturbance (terms inside the square brackets) and another accounting for the portions of the disturbance featuring systematic variation. This structure resembles closely the two components identified in our recent phenomenological study of sinusoidal adaptation data [11,12,43,44].

*Baseline drift: What is $m$ and why do we need it?*

The first term of **Equation *A1*** describes a decaying drift similar to what we observed in the data. It is interesting to note the structure of this *response* to *no-perturbation* stimulus, since the bias $m$ accounts for intrinsic properties of the visuomotor system and the term involving the stimulus $v(i)$ has been left out. To understand better its evolution with the trial number, it can be rewritten as:

$$b(n+1) = \frac{m}{1-(A-K)} + \left[G - \frac{m}{1-(A-K)}\right](A-K)^n \qquad (A2)$$

The first term of **Equation *A2*** is independent of $n$ and therefore represents an asymptotic value once all dependence on the trial number had subsided. The second term has a coefficient in front that depends on the initial condition ($G$) and, because there is no true stimulus except for that intitial value and the bias $m$, it will decay as the trial number progresses as long as $(A-K)$ is smaller than 1. In this case, it can be rewritten in terms of a *timescale* defined by the identity: $e^{-\lambda} \equiv (A-K)$. Both **Equation *A1*** and ***A2*** indicate that $(A-K)^n$ are coefficients that weigh the contribution of the stimulus to the response. This *convolution* arises naturally as a consequence of iterating the simple version of the equation (for example, that of model $KAm$ or $KAmG$). Note that, as long as $0 < (A-K) < 1$, the smaller the value of this first-trial weight, the faster its power will go to zero as the trial number $n$ increases. Thus, we define an *integration window* given by the number of trials that it takes for the weight to decrease to a size of ${}^{1}\!/_{e}$. Therefore, this integration window has size equal to $(\lambda)^{-1}$ trials. Therefore, the closer the parameter combination given by $(A-K)$ gets to 1, the smaller the value of timescale $\lambda$, and the larger the window of integration, over which collecting weighted contributions from the stimulus adds significantly to the response.

The simplest state-equation (cf. [19] ) is recovered when $A = 1$, and $m = 0$. In that case the asymptote for our adaptation gain vanishes and one would expect the baseline to remain unchanged. However, we observed a pervasive drift towards higher hypometria across all conditions in the majority of the participants. This inward drift of the baseline can be modeled assuming that $m < 0$, which could be interpreted as a systematic tendency to undershoot. As mentioned above, the sinusoidal stimulus does not contain any constant part because the disturbance is fully sinusoidal and centered at zero mean. Therefore, $m$ is required to model the drift that we observed in the adaptation gain.

This is not the only reason why the simplest version of the state equation needs to be modified. Already in the case of a disturbance that only includes a constant part $c$ (of positive or negative sign for outward or inward adaptation respectively), **Equation *A2*** will acquire an extra term proportional to that constant part, weighted by the learning rate $K$:

$$b(n+1) = \frac{m}{1-(A-K)} + \left[G - \frac{m}{1-(A-K)}\right](A-K)^n + \frac{K}{1-(A-K)}c[1-(A-K)^n\,] \qquad (A3)$$

Under the simplest state-equation ($m = 0, A = 1$) all that remains from **Equation *A3*** is $b(n+1) = c[1-(1-K)^n\,]$. Therefore, the asymptote would become $c$ predicting fully complete adaptation, which is not typically observed in paradigms that employ fixed-size or random disturbances (Herman et al. 2013). The presence of $m$ would prevent full adaptation. Lack of full completeness of adaptation could also be a consequence of a retention parameter $A < 1$. However, because the disturbance that we used does not have a constant component, even if $A < 1$, the model will not produce any drift of the baseline (cf. **Equations *A2*** and ***A3***). In experimental protocols that use fixed-step adaptation, where the disturbance has a constant component, this confound between the effects of $m$ and $A$ cannot be avoided. Hence, in the main text, we stress the importance of $m$ in our model and we will discuss possible origins and alternative interpretations for such term.

*Stimulus convolution.*

The second term in the RHS of **Equation *A1*** is a convolution of the variable part of the disturbance. For the sinusoidal disturbances used in our datasets, it produces the sinusoidal component of the oculomotor response.

$$h(n+1) = K \sum_{i=1}^{n} p(i) \sin(\omega i)\,(A-K)^{n-i}. \qquad (A4)$$

The latest experienced stimulus (at trial $n$) will be fully weighted because $i = n$. Increasing powers of $(A - K)$ will progressively attenuate subsequent older instances of the stimulus until $i$ becomes small enough so that $(A-K)^{n-i}$ becomes negligibly small. This defines a window of trials over which the stimulus contributes in a relevant manner to the oculomotor response. Note that **Equation *A4*** resembles a harmonic components expansion of a pattern with fundamental frequency given by $\omega \sim \frac{1}{T}$, where $T$ is commensurate with the intertrial interval. By design $\omega = \frac{2\pi f}{N}$, where $f$ is the frequency of the stimulus in saccades per block. This expansion of the resulting response start with a fundamental frequency determined by the inter-trial interval $T$.

*Baseline drift in models with double error-sampling ($D$).*

Next we consider, based on **Equation 14**, how the evolution of the baseline drift is expected to behave and how the timescale and window of integration change upon including a non-zero $D$. When $D = 0$, **Equation 14** gave us the first-trial weight. We obtained all weights that enter the responses in **Equation *A1*** and ***A2*** (the weights that conform the impulse response or the weights of the stimulus convolution) by raising the first-trial weight to the trial number. We extracted these relations by simply iterating **Equation 5** without $D$. When $D$ is present and non zero, an expression similar to **Equation *A2*** can be obtained, except that the term evolving with the trial number has two contributions of the form

$$\sim\frac{1}{2}\left\{\alpha(G_1,G_2)\left[(A-K)+\sqrt{(A-K)^2+4\cdot D}\right]^n+\beta(G_1,G_2)\left[(A-K)-\sqrt{(A-K)^2+4\cdot D}\right]^n\right\}, \qquad (A5)$$

where $G_1$and $G_2$ are the values of the gain in the first two trials (initial conditions). To visualize the changes, using the fitted values of the parameters we computed the corresponding first-trial weights that enter **Equation *A2*** and ***A5*** for each participant. We then took the average of these weights in each condition and ploted the sequence given by the first-trial weight raised to the trial difference with respect the current trial. In other words, we plotted the weights that would enter a stimulus convolution or, equivalently, the weights that form the impulse response for the corresponding model. The results are shown in **Fig S1** for models $KAm$ (solid lines) and $KmDG$ (dashed lines). The plots show a sizeable increase in the window of integration in model $KmDG$ with respect to that resulting for the parameters of model $KAm$, as pointed out in the **Results** section.

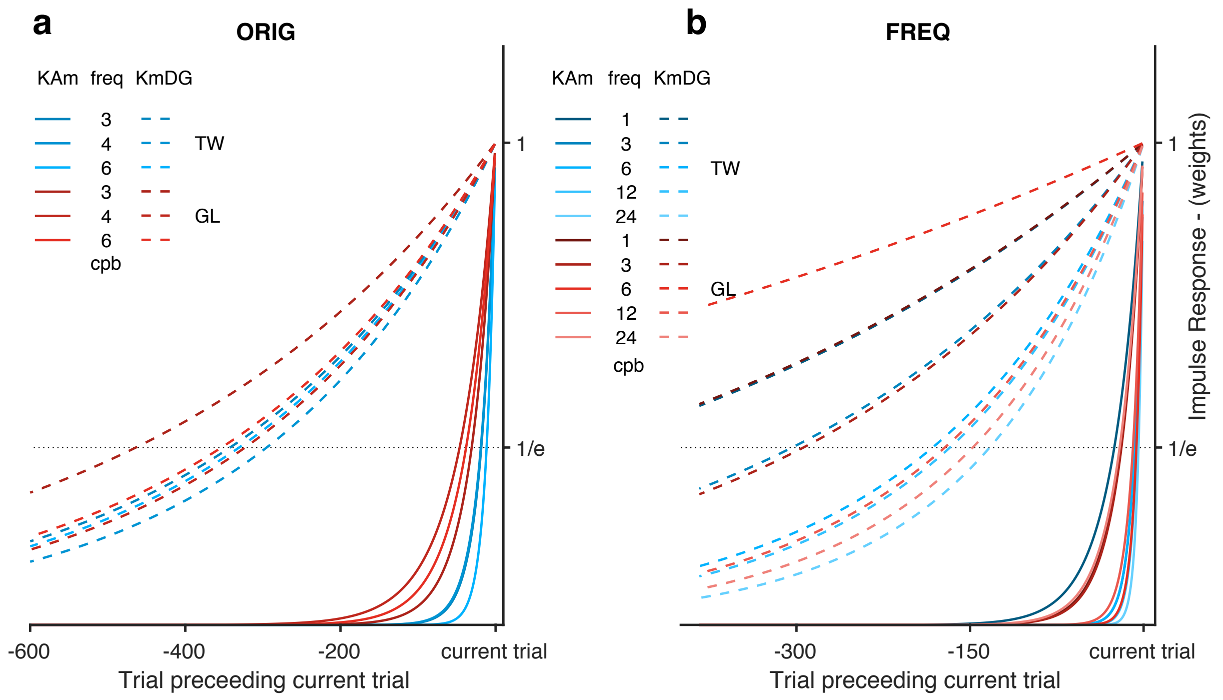


**Fig S1. Exploration of the 'window of integration'.** Weights of the convolution term in **Equation *A4*** and ***A5*** (cf. also **Equation 14** in the **Methods** section) as a function of the trial number for models $\boldsymbol{KAm}$ (solid lines) and $\boldsymbol{KmDG}$ (dashed lines), for dataset ORIG **(a)** and FREQ **(b)** analyzed in the manuscript. To determine the weights, we used the average of the timescales computed with the parameters fitted to the individual data in each condition. The number of trials that takes for the magnitude of the weights to decay to 1/e of its initial value gives an estimate of the window of integration. Including double error-sampling and the second learning rate $\boldsymbol{D}$ produces a significant increase in the window of integration with respect to the model without double error-sampling. In general, in models without $\boldsymbol{D}$ the first-trial weight always equals $\boldsymbol{A} - \boldsymbol{K}$, so that higher learning rates and smaller persistence rates result in smaller integration windows. This rigidity in the weighting of the experienced stimulus can be softened by keeping memory of, and learning from further errors in the past.
