## Supplemental Table 1 for "A generative learning model for saccade adaptation"

**Table 1. Generative parameters fitted with the best four models.**

| Model $KAm$ (ORIG) | | | | | | | |
|---|---|---|---|---|---|---|---|
| Adaptation type | Frequency | Learning rate $K$ | | Persistence rate $A$ | | Bias parameter $m$ | |
| | | ***mean*** | ***SEM*** | ***mean*** | ***SEM*** | ***mean*** | ***SEM*** |
| Two-way | 3 cpb | 0.0119 | 0.0027 | 0.9594 | 0.0110 | - 0.0137 | 0.0041 |
| | 4 cpb | 0.0109 | 0.0027 | 0.9599 | 0.0143 | - 0.0156 | 0.0056 |
| | 6 cpb | 0.0135 | 0.0039 | 0.9310 | 0.0266 | - 0.0224 | 0.0080 |
| Global | 3 cpb | 0.0052 | 0.0012 | 0.9733 | 0.0077 | - 0.0145 | 0.0048 |
| | 4 cpb | 0.0046 | 0.0007 | 0.9833 | 0.0051 | - 0.0107 | 0.0040 |
| | 6 cpb | 0.0045 | 0.0015 | 0.9786 | 0.0094 | - 0.0117 | 0.0044 |
| Model $KAm$ (FREQ) | | | | | | | |
| Two-way | 1 cpb | 0.0121 | 0.0015 | 0.9734 | 0.0046 | - 0.0128 | 0.0037 |
| | 3 cpb | 0.0254 | 0.0087 | 0.9042 | 0.0342 | - 0.0398 | 0.0204 |
| | 6 cpb | 0.0217 | 0.0052 | 0.9014 | 0.0259 | - 0.0382 | 0.0125 |
| | 12 cpb | 0.0247 | 0.0079 | 0.8703 | 0.0629 | - 0.0429 | 0.0260 |
| | 24 cpb | 0.0374 | 0.0162 | 0.8337 | 0.1151 | - 0.0450 | 0.0244 |
| Global | 1 cpb | 0.0092 | 0.0024 | 0.9595 | 0.0201 | - 0.0103 | 0.0030 |
| | 3 cpb | 0.0103 | 0.0044 | 0.9595 | 0.0190 | - 0.0224 | 0.0119 |
| | 6 cpb | 0.0139 | 0.0068 | 0.8654 | 0.0703 | - 0.0518 | 0.0272 |
| | 12 cpb | 0.0153 | 0.0062 | 0.9123 | 0.0686 | - 0.0325 | 0.0216 |
| | 24 cpb | 0.0073 | 0.0038 | 0.9600 | 0.0184 | - 0.0191 | 0.0077 |

| Model $KAmG$ (ORIG) | | | | | | | |
|---|---|---|---|---|---|---|---|
| Adaptation type | Frequency | Learning rate $K$ | | Persistence rate $A$ | | Bias parameter $m$ | |
| | | ***mean*** | ***SEM*** | ***mean*** | ***SEM*** | ***mean*** | ***SEM*** |
| Two-way | 3 cpb | 0.0087 | 0.0023 | 0.9739 | 0.0109 | - 0.0091 | 0.0039 |
| | 4 cpb | 0.0078 | 0.0020 | 0.9785 | 0.0092 | - 0.0118 | 0.0057 |
| | 6 cpb | 0.0097 | 0.0028 | 0.9463 | 0.0258 | - 0.0152 | 0.0070 |
| Global | 3 cpb | 0.0045 | 0.0012 | 0.9832 | 0.0067 | - 0.0087 | 0.0031 |
| | 4 cpb | 0.0038 | 0.0008 | 0.9932 | 0.0026 | - 0.0045 | 0.0015 |
| | 6 cpb | 0.0049 | 0.0016 | 0.9794 | 0.0094 | - 0.0101 | 0.0041 |
| Model $KAmG$ (FREQ) | | | | | | | |
| Two-way | 1 cpb | 0.0103 | 0.0020 | 0.9811 | 0.0051 | - 0.0081 | 0.0036 |
| | 3 cpb | 0.0234 | 0.0097 | 0.8725 | 0.0842 | - 0.0241 | 0.0090 |
| | 6 cpb | 0.0132 | 0.0048 | 0.9638 | 0.0211 | - 0.0147 | 0.0069 |
| | 12 cpb | 0.0144 | 0.0052 | 0.9506 | 0.0365 | - 0.0392 | 0.0262 |
| | 24 cpb | 0.0334 | 0.0158 | 0.8293 | 0.1300 | - 0.0645 | 0.0382 |
| Global | 1 cpb | 0.0056 | 0.0006 | 0.9838 | 0.0032 | - 0.0066 | 0.0021 |
| | 3 cpb | 0.0056 | 0.0010 | 0.9819 | 0.0068 | - 0.0096 | 0.0032 |
| | 6 cpb | 0.0024 | 0.0023 | 1.0010 | 0.0121 | - 0.0020 | 0.0065 |
| | 12 cpb | 0.0139 | 0.0067 | 0.9195 | 0.0735 | - 0.0286 | 0.0230 |
| | 24 cpb | 0.0045 | 0.0036 | 0.9993 | 0.0044 | - 0.0026 | 0.0010 |

| Model $KmDG$ (ORIG) | | | | | | | |
|---|---|---|---|---|---|---|---|
| Adaptation type | Frequency | Learning rate $K$ | | Learning rate $D$ | | Bias parameter $m$ | |
| | | ***mean*** | ***SEM*** | ***mean*** | ***SEM*** | ***mean*** | ***SEM*** |
| Two-way | 3 cpb | 0.1575 | 0.0168 | - 0.1541 | 0.0167 | -0.0016 | 0.0003 |
| | 4 cpb | 0.1351 | 0.0137 | - 0.1313 | 0.0136 | - 0.0017 | 0.0003 |
| | 6 cpb | 0.1267 | 0.0173 | - 0.1232 | 0.0170 | - 0.0015 | 0.0003 |
| Global | 3 cpb | 0.0667 | 0.0177 | - 0.0644 | 0.0176 | - 0.0013 | 0.0002 |
| | 4 cpb | 0.0448 | 0.0109 | - 0.0416 | 0.0107 | - 0.0018 | 0.0005 |
| | 6 cpb | 0.0349 | 0.0179 | - 0.0320 | 0.0175 | - 0.0012 | 0.0007 |
| Model $KmDG$ (FREQ) | | | | | | | |
| Two-way | 1 cpb | 0.2235 | 0.0275 | - 0.2210 | 0.0273 | - 0.0013 | 0.0002 |
| | 3 cpb | 0.1568 | 0.0308 | - 0.1530 | 0.0308 | - 0.0017 | 0.0004 |
| | 6 cpb | 0.1233 | 0.0169 | - 0.1171 | 0.0165 | - 0.0026 | 0.0004 |
| | 12 cpb | 0.0925 | 0.0101 | - 0.0861 | 0.0107 | - 0.0031 | 0.0007 |
| | 24 cpb | 0.0831 | 0.0168 | - 0.0751 | 0.0166 | - 0.0035 | 0.0010 |
| Global | 1 cpb | 0.1018 | 0.0167 | - 0.0996 | 0.0168 | - 0.0012 | 0.0002 |
| | 3 cpb | 0.0733 | 0.0151 | - 0.0697 | 0.0152 | - 0.0019 | 0.0003 |
| | 6 cpb | 0.0521 | 0.0134 | - 0.0509 | 0.0132 | - 0.0009 | 0.0007 |
| | 12 cpb | 0.0310 | 0.0110 | - 0.0251 | 0.0112 | - 0.0019 | 0.0005 |
| | 24 cpb | 0.0340 | 0.0095 | - 0.0271 | 0.0086 | - 0.0029 | 0.0009 |

| Model $KAmDG$ (ORIG) | | | | | | | | | |
|---|---|---|---|---|---|---|---|---|---|
| Adaptation type | Frequency | Learning rate $K$ | | Learning rate $D$ | | Persistence rate $A$ | | Bias parameter $m$ | |
| | | ***Mean*** | ***SEM*** | ***Mean*** | ***SEM*** | ***mean*** | ***SEM*** | ***mean*** | ***SEM*** |
| Two-way | 3 cpb | 0.1552 | 0.0170 | - 0.1519 | 0.0170 | 0.9994 | 0.0013 | - 0.0020 | 0.0006 |
| | 4 cpb | 0.1199 | 0.0206 | - 0.1152 | 0.0215 | 0.9950 | 0.0063 | - 0.0029 | 0.0014 |
| | 6 cpb | 0.1255 | 0.0178 | - 0.1216 | 0.0174 | 0.9973 | 0.0017 | - 0.0022 | 0.0004 |
| Global | 3 cpb | 0.0784 | 0.0197 | - 0.0761 | 0.0196 | 1.0014 | 0.0033 | - 0.0014 | 0.0015 |
| | 4 cpb | 0.0254 | 0.0147 | - 0.0215 | 0.0145 | 0.9892 | 0.0057 | - 0.0081 | 0.0039 |
| | 6 cpb | 0.0328 | 0.0166 | - 0.0294 | 0.0161 | 0.9932 | 0.0032 | - 0.0042 | 0.0011 |
| Model $KAmDG$ (FREQ) | | | | | | | | | |
| Two-way | 1 cpb | 0.1642 | 0.0566 | - 0.1588 | 0.0583 | 0.9914 | 0.0063 | - 0.0058 | 0.0034 |
| | 3 cpb | 0.1338 | 0.0263 | - 0.1255 | 0.0257 | 0.9828 | 0.0086 | - 0.0108 | 0.0061 |
| | 6 cpb | 0.1177 | 0.0222 | - 0.1099 | 0.0229 | 0.9902 | 0.0163 | - 0.0086 | 0.0067 |
| | 12 cpb | 0.0912 | 0.0105 | - 0.0835 | 0.0103 | 0.9917 | 0.0077 | - 0.0093 | 0.0052 |
| | 24 cpb | 0.0850 | 0.0171 | - 0.0735 | 0.0161 | 0.9976 | 0.0061 | - 0.0069 | 0.0029 |
| Global | 1 cpb | 0.0518 | 0.0567 | - 0.0476 | 0.0581 | 0.9905 | 0.0085 | - 0.0067 | 0.0035 |
| | 3 cpb | 0.0505 | 0.0278 | - 0.0452 | 0.0294 | 0.9883 | 0.0104 | - 0.0095 | 0.0060 |
| | 6 cpb | 0.0568 | 0.0159 | - 0.0547 | 0.0154 | 0.9934 | 0.0086 | - 0.0026 | 0.0033 |
| | 12 cpb | 0.0319 | 0.0112 | - 0.0254 | 0.0114 | 1.0017 | 0.0030 | - 0.0015 | 0.0010 |
| | 24 cpb | 0.0318 | 0.0094 | - 0.0290 | 0.0085 | 0.9978 | 0.0045 | - 0.0025 | 0.0010 |

Model name and dataset are shown at the top. The corresponding datasets can also be identified by the stimulus frequencies tested: ORIG: 3, 4 and 6cpb. FREQ: 1, 3, 6, 12 and 24cpb. The initial condition parameter is not reported.
