## Supplemental Table 2 for "A generative learning model for saccade adaptation"

**Table 2. ANOVA results on the generative parameters fitted with the best four models.**

| Dataset FREQ | | $K$ | | $A$ | | $m$ | | $D$ | | $G$ | | $\kappa = K + D$ | | $\eta = K - D$ | |
|---|---|---|---|---|---|---|---|---|---|---|---|---|---|---|---|
| Model | Regressor | $F$ | $p > F$ | $F$ | $p > F$ | $F$ | $p > F$ | $F$ | $p > F$ | $F$ | $p > F$ | $F$ | $p > F$ | $F$ | $p > F$ |
| $KAm$ | Freq | 0.89 | 0.47 | 0.89 | 0.48 | 1.01 | 0.41 | | | | | | | | |
| | Type | 8.07 | 0.01 | 1.96 | 0.19 | 1.22 | 0.29 | | | | | | | | |
| | Subj | 1.84 | 0.22 | 2.85 | 0.13 | 2.96 | 0.10 | N/A | | N/A | | N/A | | N/A | |
| | Freq*Type | 1.10 | 0.37 | 0.75 | 0.56 | 0.41 | 0.80 | | | | | | | | |
| | Freq*Subj | 0.86 | 0.71 | 1.00 | 0.49 | 1.11 | 0.36 | | | | | | | | |
| | Type*Subj | 1.05 | 0.42 | 0.57 | 0.85 | 0.54 | 0.88 | | | | | | | | |
| $KAmG$ | Freq | 1.18 | 0.33 | 0.68 | 0.61 | 1.35 | 0.27 | | | 0.10 | 0.98 | | | | |
| | Type | 8.57 | 0.01 | 2.70 | 0.13 | 2.71 | 0.13 | | | 0.01 | 0.93 | | | | |
| | Subj | 0.90 | 0.59 | 0.75 | 0.69 | 0.78 | 0.67 | | | 2.34 | 0.05 | | | | |
| | Freq*Type | 1.30 | 0.28 | 1.16 | 0.34 | 1.18 | 0.33 | N/A | | 1.68 | 0.17 | N/A | | N/A | |
| | Freq*Subj | 0.80 | 0.78 | 1.04 | 0.44 | 1.05 | 0.43 | | | 1.75 | 0.03 | | | | |
| | Type*Subj | 0.95 | 0.50 | 1.07 | 0.41 | 1.58 | 0.13 | | | 0.84 | 0.61 | | | | |
| $KmDG$ | Freq | 21.38 | 0.00 | | | 4.60 | < 0.01 | 23.16 | 0.00 | 1.03 | 0.40 | 5.97 | < 0.001 | 22.31 | 0.00 |
| | Type | 35.12 | < 0.01 | | | 3.33 | 0.09 | 34.69 | < 0.001 | 0.00 | 0.99 | 1.63 | 0.23 | 34.97 | < 0.001 |
| | Subj | 2.81 | 0.05 | | | 2.45 | 0.10 | 2.84 | 0.05 | 13.62 | < 0.001 | 0.90 | 0.59 | 2.83 | 0.05 |
| | Freq*Type | 2.16 | 0.09 | N/A | | 1.13 | 0.35 | 2.29 | 0.07 | 0.83 | 0.51 | 0.92 | 0.46 | 2.23 | 0.09 |
| | Freq*Subj | 0.97 | 0.54 | | | 0.89 | 0.65 | 0.99 | 0.51 | 1.31 | 0.18 | 0.65 | 0.93 | 0.98 | 0.52 |
| | Type*Subj | 2.38 | 0.02 | | | 1.33 | 0.24 | 2.36 | 0.02 | 0.62 | 0.82 | 1.33 | 0.23 | 2.37 | 0.02 |
| $KAmDG$ | Freq | 1.24 | 0.31 | 0.72 | 0.58 | 0.57 | 0.69 | 1.30 | 0.28 | 0.12 | 0.98 | 0.64 | 0.64 | 1.27 | 0.30 |
| | Type | 15.81 | < 0.01 | 0.59 | 0.46 | 2.00 | 0.18 | 14.30 | < 0.01 | 0.01 | 0.94 | 5.17 | 0.04 | 15.08 | < 0.01 |
| | Subj | 1.81 | 0.25 | 1.81 | 0.27 | 1.68 | 0.22 | 1.96 | 0.24 | 3.65 | < 0.01 | 0.62 | 0.80 | 1.89 | 0.25 |
| | Freq*Type | 0.34 | 0.85 | 0.13 | 0.97 | 0.39 | 0.81 | 0.38 | 0.82 | 2.03 | 0.10 | 1.85 | 0.14 | 0.36 | 0.84 |
| | Freq*Subj | 0.82 | 0.76 | 0.89 | 0.65 | 1.04 | 0.45 | 0.80 | 0.78 | 2.30 | < 0.01 | 1.48 | 0.09 | 0.81 | 0.77 |
| | Type*Subj | 0.98 | 0.48 | 0.74 | 0.70 | 1.12 | 0.36 | 0.91 | 0.54 | 2.02 | 0.04 | 2.73 | < 0.01 | 0.94 | 0.52 |

| Dataset ORIG | | $K$ | | $A$ | | $m$ | | $D$ | | $G$ | | $\kappa = K + D$ | | $\eta = K - D$ | |
|---|---|---|---|---|---|---|---|---|---|---|---|---|---|---|---|
| Model | Regressor | $F$ | $p > F$ | $F$ | $p > F$ | $F$ | $p > F$ | $F$ | $p > F$ | $F$ | $p > F$ | $F$ | $p > F$ | $F$ | $p > F$ |
| $KAm$ | Freq | 0.23 | 0.80 | 0.74 | 0.49 | 0.36 | 0.70 | N/A | | N/A | | N/A | | N/A | |
| | Type | 12.64 | < 0.01 | 8.09 | 0.02 | 2.18 | 0.17 | | | | | | | | |
| | Subj | 1.58 | 0.30 | 1.17 | 0.42 | 2.79 | 0.23 | | | | | | | | |
| | Freq*Type | 0.26 | 0.77 | 1.09 | 0.36 | 0.70 | 0.51 | | | | | | | | |
| | Freq*Subj | 0.90 | 0.59 | 1.43 | 0.23 | 0.97 | 0.53 | | | | | | | | |
| | Type*Subj | 1.64 | 0.18 | 1.07 | 0.43 | 0.70 | 0.70 | | | | | | | | |
| $KAmG$ | Freq | 0.39 | 0.68 | 1.45 | 0.26 | 0.5 | 0.62 | N/A | | 0.82 | 0.46 | N/A | | N/A | |
| | Type | 12.66 | < 0.01 | 6.61 | 0.03 | 5.77 | 0.04 | | | 4.97 | 0.05 | | | | |
| | Subj | 2.13 | 0.10 | 1.49 | 0.25 | 2.35 | 0.16 | | | 3.10 | 0.14 | | | | |
| | Freq*Type | 0.05 | 0.95 | 1.01 | 0.38 | 0.49 | 0.62 | | | 0.51 | 0.61 | | | | |
| | Freq*Subj | 2.20 | 0.05 | 2.46 | 0.03 | 1.86 | 0.10 | | | 0.64 | 0.82 | | | | |
| | Type*Subj | 1.58 | 0.20 | 1.06 | 0.43 | 0.37 | 0.93 | | | 1.56 | 0.20 | | | | |
| $KmDG$ | Freq | 5.31 | 0.02 | N/A | | 0.62 | 0.55 | 5.24 | 0.02 | 0.37 | 0.70 | 0.64 | 0.54 | 5.27 | 0.02 |
| | Type | 36.94 | < 0.001 | | | 0.35 | 0.57 | 37.57 | < 0.001 | 6.07 | 0.04 | 2.45 | 0.15 | 37.26 | < 0.001 |
| | Subj | 1.91 | 0.17 | | | 1.40 | 0.36 | 1.87 | 0.17 | 4.46 | 0.13 | 1.10 | 0.45 | 1.89 | 0.17 |
| | Freq*Type | 0.00 | 1.00 | | | 0.14 | 0.87 | 0.01 | 0.99 | 0.09 | 0.91 | 0.21 | 0.81 | 0.00 | 1.00 |
| | Freq*Subj | 1.25 | 0.32 | | | 1.10 | 0.42 | 1.35 | 0.27 | 0.64 | 0.82 | 1.54 | 0.18 | 1.30 | 0.29 |
| | Type*Subj | 4.31 | < 0.01 | | | 1.09 | 0.41 | 4.29 | < 0.01 | 1.20 | 0.35 | 1.24 | 0.33 | 4.30 | < 0.01 |
| $KAmDG$ | Freq | 3.82 | 0.04 | 1.62 | 0.23 | 2.26 | 0.13 | 3.82 | 0.04 | 0.07 | 0.94 | 1.88 | 0.18 | 3.82 | 0.04 |
| | Type | 30.75 | < 0.001 | 1.10 | 0.32 | 2.09 | 0.18 | 31.03 | < 0.001 | 3.65 | 0.09 | 2.19 | 0.17 | 30.90 | < 0.001 |
| | Subj | 0.99 | 0.48 | 0.88 | 0.56 | 1.01 | 0.49 | 0.94 | 0.51 | 3.79 | 0.11 | 1.71 | 0.31 | 0.97 | 0.50 |
| | Freq*Type | 0.64 | 0.54 | 1.02 | 0.38 | 2.01 | 0.16 | 0.70 | 0.51 | 0.25 | 0.78 | 0.09 | 0.92 | 0.67 | 0.52 |
| | Freq*Subj | 4.07 | < 0.01 | 2.61 | 0.02 | 1.60 | 0.16 | 4.47 | < 0.01 | 0.69 | 0.78 | 1.14 | 0.39 | 4.27 | < 0.01 |
| | Type*Subj | 5.15 | < 0.01 | 1.09 | 0.42 | 1.69 | 0.16 | 5.19 | < 0.01 | 1.42 | 0.25 | 1.74 | 0.67 | 5.18 | < 0.01 |

Repeated-measures ANOVA (2 X 3 on data from ORIG; 2 X 5 on data from FREQ) with factors type of adaptation and stimulus frequency was run on each of the four best models. Model name is shown at the side of the table and parameter names are on the top. The dataset is indicated in the cell at the upper left corner next to the parameter names. Highlights indicate the cases where the corresponding factor reached significance level.
